# 3D single-cell shape analysis using geometric deep learning

**DOI:** 10.1101/2022.06.17.496550

**Authors:** Matt De Vries, Lucas Dent, Nathan Curry, Leo Rowe-Brown, Vicky Bousgouni, Adam Tyson, Christopher Dunsby, Chris Bakal

## Abstract

Aberrations in 3D cell morphogenesis are linked to diseases such as cancer. Yet there is little systems-level understanding of cell shape determination in 3D, largely because there is a paucity of data-driven methods to quantify and describe 3D cell shapes. We have addressed this need using unsupervised geometric deep learning to learn shape representations of over 95,000 melanoma cells imaged by 3D high-throughput light-sheet microscopy. We used a dynamic graph convolutional foldingnet autoencoder with improved deep embedded clustering to simultaneously learn lower-dimensional representations and classes of 3D cell shapes. We describe a landscape of 3D cell morphology using deep learning-derived 3D quantitative morphological signatures (3DQMS) across different substrate geometries, following treatment by different clinically relevant small molecules and systematic gene depletion in high-throughput. By data integration, we predict modes of action for different small molecules providing mechanistic insights and blueprints for biological re-engineering. Finally, we provide explainability and interpretability for deep learning models.

## Introduction

Advances in high throughput light-sheet microscopy [1, 2] have made it possible to image large numbers of cells in three dimensions (3D) with sub-cellular resolution. This development has led to a growing need for accurate and quantitative 3D representations of biological structures. Robust representation of the shape of 3D objects is a fundamental goal in computer vision, and there has been rapid progress in this domain. Deep learning (DL)-based shape analysis methods have been built on large-scale labelled datasets of 3D models of everyday objects [3, 4, 5]. State-of-the-art methods have analysed these 3D objects as either 3D binary voxels or point clouds for tasks of classification [6, 7, 8, 9], and unsupervised representation learning [10, 11, 12, 13, 14]. This has led to in-depth research into DL architectures which explore local geometric information using convolution, graph, and attention mechanisms on point clouds for improvements in 3D shape analysis. Geometric deep learning (GDL) involves techniques that attempt to generalise deep learning methods to non-Euclidean data, such as graphs or manifolds [15]. For example, graph-based autoencoders utilise GDL to learn feature representations of 3D shapes [16, 17, 18].

Motivated by the need to extract information embedded in 3D cell shapes and by the effectiveness of GDL approaches in generating 3D shape representations of everyday objects, we have used GDL to learn the landscape, or latent space, of 3D cell morphogenesis. To understand the environmental drivers of 3D shape determination, we use our GDL features to quantify 3D cell shape as melanoma cells explore hydrogel matrices of different geometries. The biological mechanism underpinning 3D morphogenesis was investigated using GDL to quantify the effect of clinically relevant small molecules on exploring latent “shape” space in an explainable and interpretable fashion. We also demonstrate the utility of our methods for high-throughput genetics by using them to analyse images acquired following systematic gene depletion. DL-derived 3D quantitative morphological signatures (3DQMS) allowed us to describe new functions and predict structural features for 168 proteins in an automated manner. Finally, we integrated independently derived datasets of 3D cell shape following small-molecule and genetic perturbation to characterise biological pathways targeted by small-molecules. Taken together, this work demonstrates that GDL-based 3D shape analysis of single cells can be used for high-throughput explainable genetics and the design of approaches to re-engineer morphogenesis for therapeutic benefit. Furthermore, we have packaged all of our methods in an easy-to-use Python package^4^.

## Results

### Data acquisition and processing

We used oblique plane microscopy (OPM) [1, 2] to perform rapid light-sheet imaging of more than 65 000 (65 500) WM266.4 metastatic melanoma cells embedded in tissue-like collagen matrices (Figure 1 A). Cells were physically embedded in an environment that spanned from mechanically rigid and flat/2D (proximal to the coverslip) to mechanically soft and 3D (distal to the coverslip). Coverslip height was identified using a *z* profile of the nucleus intensity as described in [19]. We also treated with clinically relevant inhibitors of the cytoskeleton and inhibitors of cell signalling pathways to sample a wide range of the possible shapes that cells can explore (see Methods and Supplementary Tables 1 and 2).

**Figure 1:**
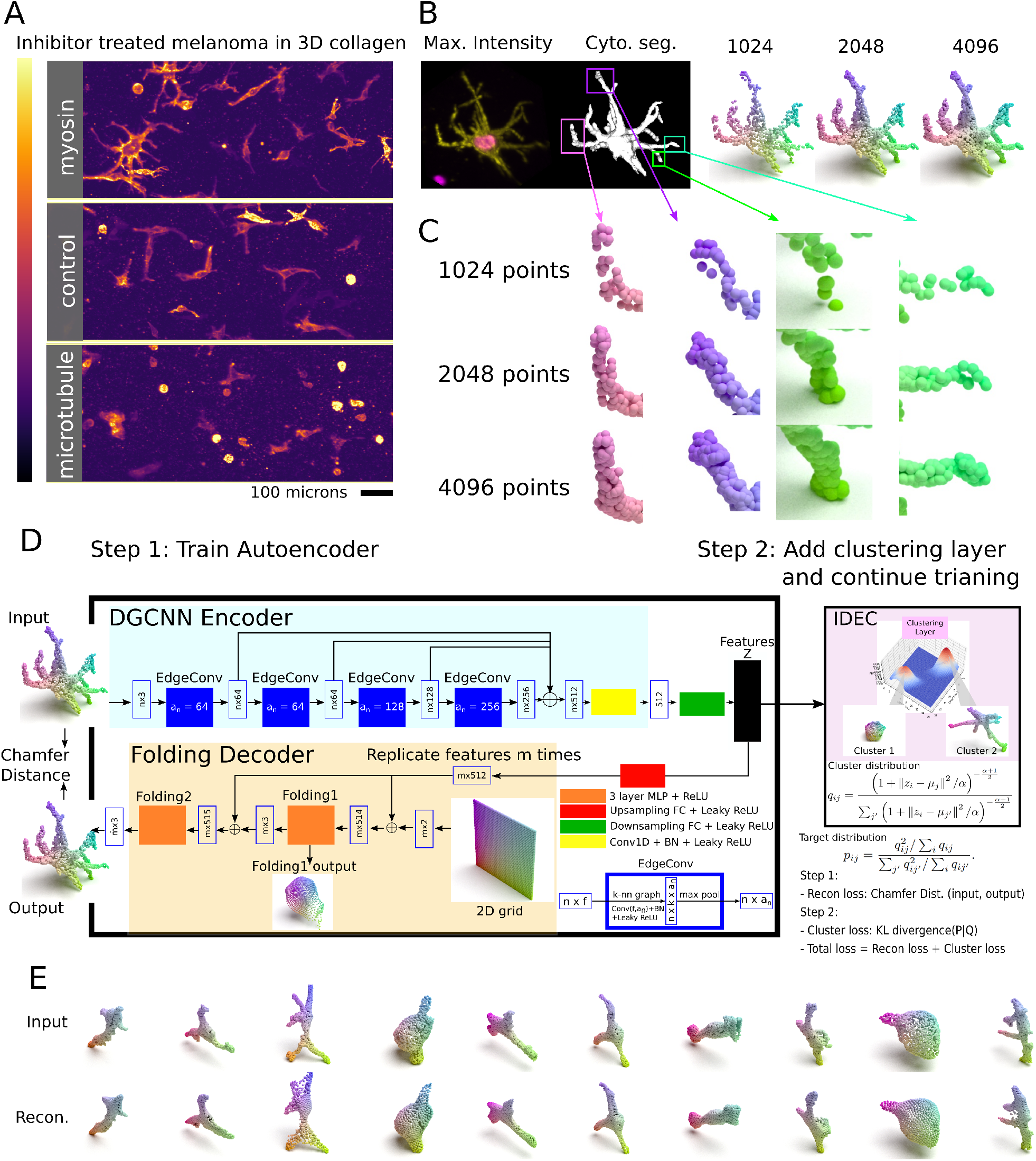
Single cell 3D shape analysis using geometric deep learning. (A) Single cells embedded in a collagen matrix were treated with a variety of small molecule drugs and imaged in 3D in high-throughput by stage scanning oblique plane microscopy (ssOPM). Maximum intensity projections (XY view) of cells treated with DMSO (control), Blebbistatin (myosin) and Nocodazole (microtubule) are shown. Intensity is the grayscale intensity of the CAAX-GFP membrane marker. (B) The cells were labelled with transgenes expressing Histone-2B (which labels the nucleus) and CAAX-GFP (which labels cell membranes). Nucleus and cell segmentation masks (3D mask) are shown for a single cell (Cyto. seg.). We used the marching cubes algorithm to create a mesh object of vertices, faces, and normals and then sampled 1024, 2048, and 4096 points from this mesh object to create a point cloud for each cell cytoplasm and nucleus. (C) A sampling density of 2046 points was sufficient enough to represent the shapes. (D) Our method is trained in two steps. Step 1 is to train the DFN autoencoder to learn a feature vector. DFN comprises a DGCNN encoder (light blue) and a FoldingNet decoder (light orange). This autoencoder is trained by minimising the Chamfer Distance between the input and output point cloud. Step 2 is to add the clustering layer (light pink) and continue training. This step refines the autoencoder weights and learns classes. A Cluster loss is added in the second step. (E) Example input and reconstructed 3D point clouds of melanoma cells.

Cells and nuclei were segmented using different segmentation methods serially, including active contours and Otsu’s thresholding techniques (see Methods). 3D voxel masks were then converted to mesh objects (Figure 1 B Cyto. seg.) using marching cubes [20], and points were sampled from the surfaces of each mesh object to give a point cloud representation of each cell and nucleus (see Methods). We tested a range of point sampling densities to ensure that points along complex cellular protrusions were sufficiently captured (Figure 1 C). Point clouds were pre-processed by mean centring and scaling to assist in training our GDL model.

### An automated method to profile cell shape in 3D using geometric deep learning

To create an automated method to profile single-cell shapes in 3D, we used GDL models (Figure 1 D). The model described in this paper has two focuses. First, we used a graph-based autoencoder to learn shape features in a self-supervised manner (Step 1 in Figure 1 D). Here, we adapted a FoldingNet [17] autoencoder by using a Dynamic Graph Convolutional Neural Network (DGCNN) [6] encoder to give a Dynamic FoldingNet (DFN). Second, we simultaneously learned shape features and classes by adding a clustering layer (Step 2 in Figure 1 D). The clustering layer learns the shape modes based on the features from the autoencoder. This second part is known as improved deep embedded clustering (IDEC) [21]. We use the clustering layer’s output to calculate a cell’s shape signature based on its similarity to the shape modes in the dataset. Initially, we report on the DFN without the additional clustering layer and then add the clustering layer in later sections. Figure 1 E shows example input and reconstructed point clouds for a trained DFN autoencoder showing the ability of the autoencoder to reconstruct complex shapes from 128 features.

### Exploring the cell shape landscape of a metastatic melanoma cell line

Our trained DFN model without the clustering layer was first used to extract 128 3D DL shape features from all cells in our dataset. We visualised the extent of the cell shape landscape by performing UMAP [22] on the extracted features. We sampled two-dimensional locations along this projection and plotted rendered images of cell masks which generated the features projected in UMAP space (Figure 2 A). The first dimension of the UMAP appears to incorporate size and eccentricity, with smaller and rounder cells having low values for this dimension and larger and more eccentric cells having high values (see Supplementary Figure 1). After visualising the extent of the cell shape landscape, we looked at the distribution of cells within this landscape. We estimated the probability density function using kernel density estimation of the UMAP embedding (Figure 2 B).

**Figure 2:**
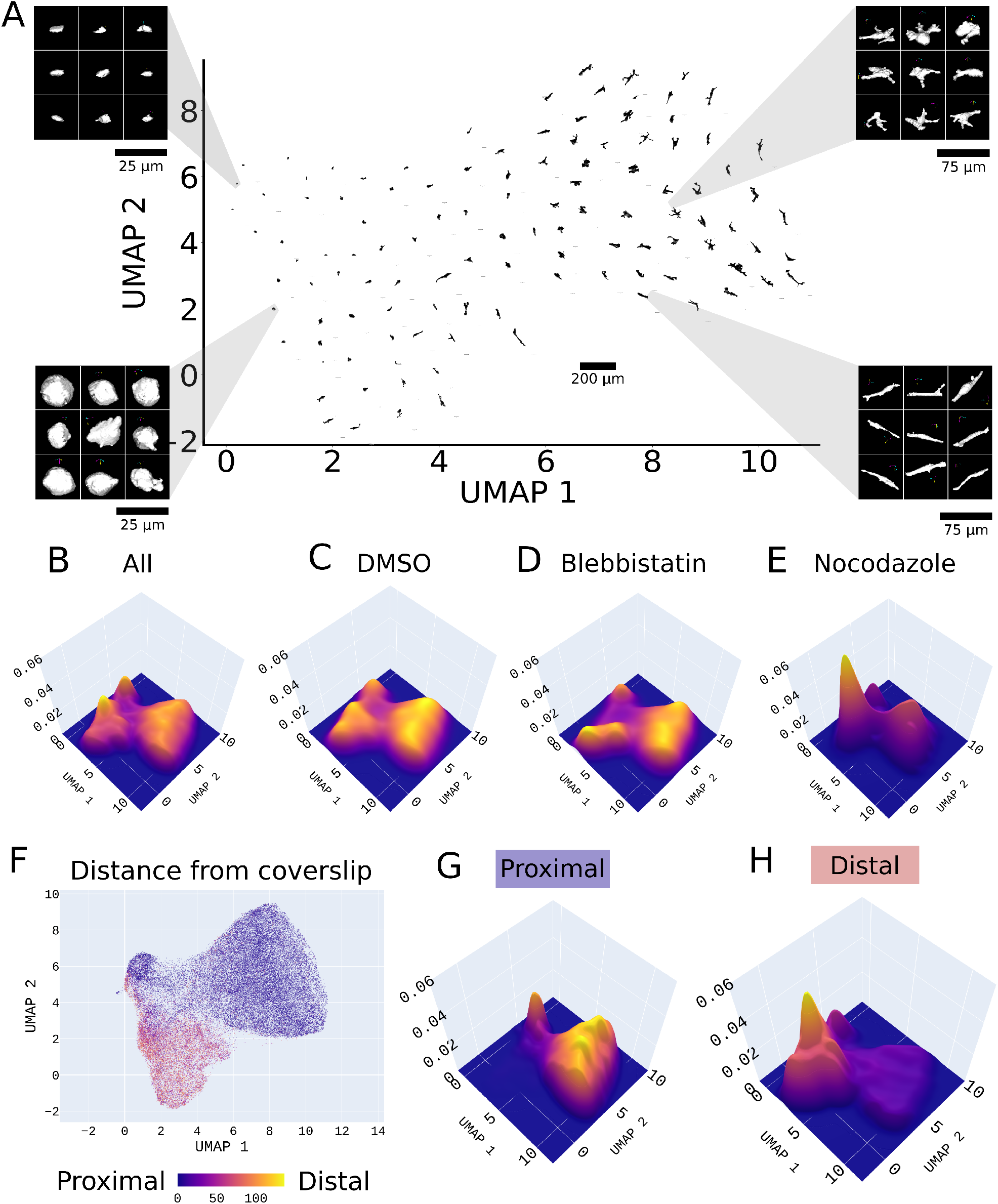
Dynamic graph convolutional foldingnet reveals the cell shape landscape in a cancer cell line. We performed UMAP on the 128 features extracted using the DFN without adding the clustering layer. (A) Rendered sample images of cell masks are plotted in UMAP. (B-E) The probability density of the UMAP was estimated using kernel density estimation. DMSO-treated cells (C) had different UMAP distributions than Blebbistatin (D) treated cells and Nocodazole (E) treated cells. (F) The points on the UMAP were coloured by their distance relative to the coverslip. The microenvironment impacts the ability of cells to take on different shapes. (G) Cells on the coverslip tend to be flatter and more spread out. (H) Cells embedded in collagen (further away from the coverslip) are more round.

A subset of cells in our dataset had been treated with small molecule inhibitors, including inhibitors of myosin (Blebbistatin) and microtubule polymerization (Nocodazole). We visually assessed the effect of these on cell shapes in 3D. The cell shape feature landscape demonstrated that cells treated with different inhibitors cause different distributions of embedding (which represent shape distributions). We observed that control (DMSO) cells could take a wide range of shapes in the cell shape space (Figure 2 C), whereas the treated cells are confined to specific areas of the shape space. For example, Blebbistatin (myosin inhibitor) treated cells (Figure 2 D) are confined to more protrusive and eccentric shapes, and cells treated with Nocodazole (microtubule polymerisation inhibitor) are confined to rounder-like cells with few protrusive cells (Figure 2 E). The effect of treatments does not create novel geometries but rather shifts the distribution of features within a population. See Supplementary Figure 2 A-J for all treatments.

The cells in our dataset are embedded in a 3D collagen matrix at various depths, with some adhering to the 2D surface of the plate itself. Thus cells encounter matrices with different geometries and stiffness. To understand how differences in the physical environment influence cell shape, we visualised our UMAP space based on the depth in the collagen. We coloured each embedding location in UMAP space according to the cell’s distance from the coverslip (Figure 2 F). This showed that cells close to the rigid coverslip (proximal to the coverslip) and cells embedded in collagen (distal to the coverslip) are separated in shape space and that the geometry and physical properties of the microenvironment have a major influence on cell shape. Following this observation, we handled cells in different depths in the collagen as being in different environments. We labelled cells within 7 *µm* of the coverslip as ‘Proximal’ and cells greater than 7 *µm* as ‘Distal’. This cutoff of 7 *µm* ensures that most cells labelled as ‘Proximal’ will be touching the coverslip, and most cells labelled as ‘Distal’ will not. The distribution of cells proximal to the coverslip (Figure 2 G) was much different than cells distal to the coverslip (Figure 2 H), with most cells in the distal setting taking on shapes which are almost non-existent in the proximal setting.

### Learned shape features enable downstream classification tasks

We explored the possibility of distinguishing between cells with different small molecule inhibitor treatments based purely on cytoplasm and nucleus shape features. We used the trained DFN autoencoder without adding the clustering layer to extract features from each cell and nucleus. These features were then used to train a support vector machine (SVM) to predict the small molecule inhibitor. We performed 10-fold cross-validation for each experiment and reported mean balanced accuracies to account for differences in class sizes. We report on our best-performing (in terms of classification accuracy) set of features.

We tested the prediction of all of our treatments in a pairwise fashion using the combination of cell and nucleus shape features (Figure 3 A and B). Due to the strong influence of the environment on shape, we made these predictions separately for proximal cells (Figure 3 A) and distal cells (Figure 3 B). We show both environments together in the supplementary material. In the proximal setting, we most accurately classified cells treated with Nocodazole, Blebbistatin and H1152 (a Rho-associated kinase (ROCK) inhibitor) from other cells with accuracies of 80%, 72.6% and 68.3% on average for these treatments in the proximal and accuracies of than 75.5%, 71.8% and 66.8% in the distal setting, respectively. Interestingly, although Binimetinib (a mitogen-activated protein kinase kinase (MEK) 1/2 inhibitor) is thought to act primarily on growth signalling rather than cell geometry, we could classify cells treated with Binimetinib from other treatments with an average accuracy of 66.6% in the proximal setting. Cells treated with Binimetinib were distinguishable from ‘No Treatment’ and DMSO, with accuracies greater than 62% in the proximal setting. The classification accuracy of Binimetinib vs all other treatments was, on average, 3.9% greater in the proximal setting (two-sided paired t-test statistic = 3.84 with p-value=0.005) and on average 4.4% greater in the proximal setting for Nocodazole vs all other treatments (two-sided paired t-test statistic of 8.04 with a p-value=4.21 × 10^−5^) (Figure 3 C). This suggests that mitogen-activated protein kinase (MAPK) signalling (inhibited by Binimetinib) and microtubule polymerisation (inhibited by Nocodazole) contribute more to cell morphogenesis in the mechanically rigid environment close to the coverslip compared to the distal environment.We also tested cytoplasm features and nuclear features alone to assess the effect on classification accuracy. Based on knowledge of the different effects on cell shape of Blebbistatin and Nocodazole, we show this in Figure 3 D (we also show this for all treatments in the Supplementary Figure 3). We found the greatest prediction accuracy resulted from combining both cytoplasmic and nuclear shape features. This difference in accuracy between combined and individual cell-component data sets suggests that not only are the shapes of the cell and nucleus affected by these treatments but so too is the relationship between them. We could also classify whether a cell was less or greater than 7 *µm* from the coverslip using cell and nucleus features with an accuracy of 85.29% (std = 1%). This demonstrates that cell shape is dependent on the environment. We saw, on average, that using GDL features to predict treatment outperformed classical features of cell geometry, including size, volume, and sphericity, by 1.02% in the distal setting and 3.08% in the proximal setting (see Supplementary Table 3 and Supplementary Figure 3).

**Figure 3:**
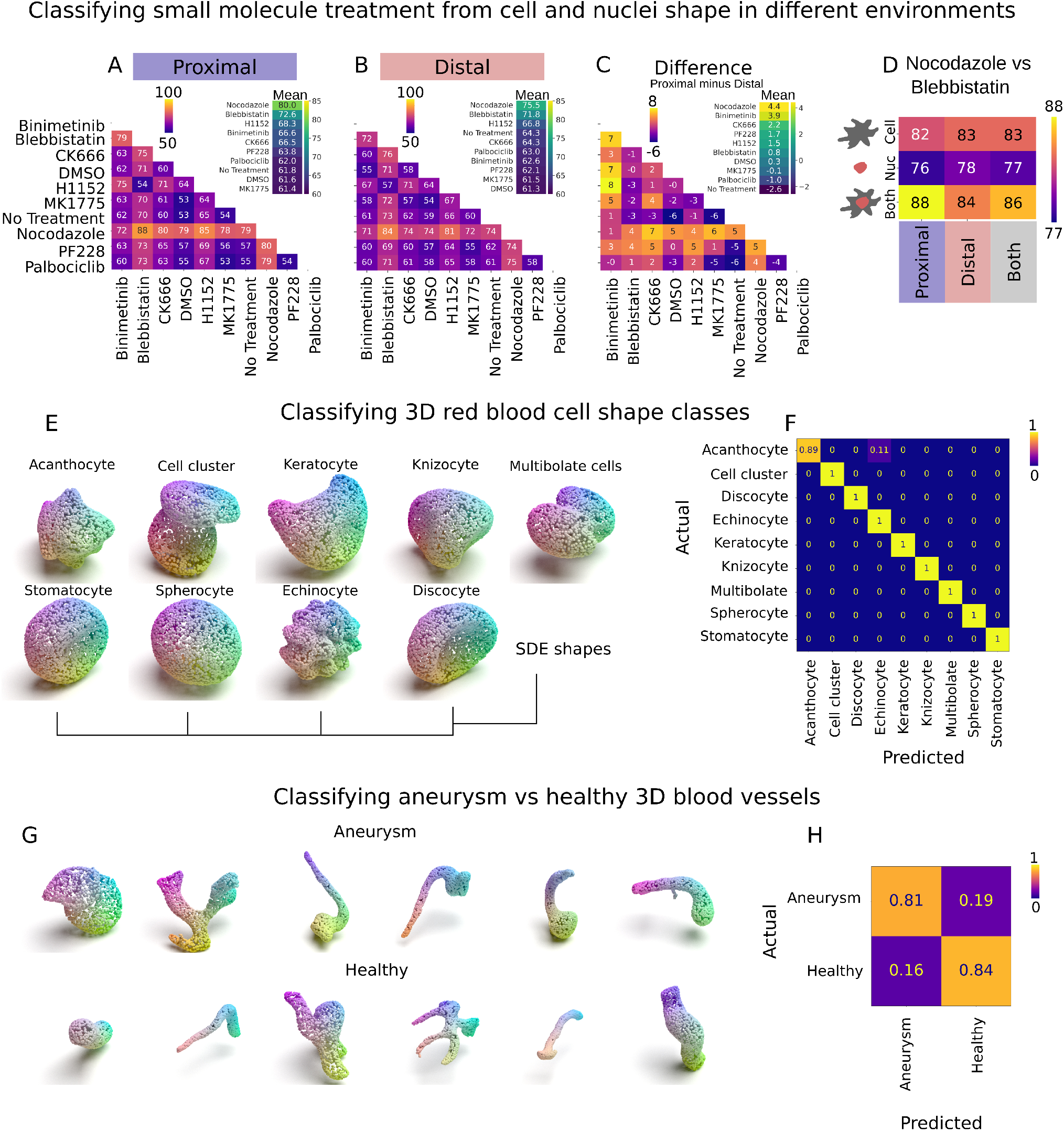
Downstream classification tasks of features extracted using DFN. (A-B) We used an SVM to predict treatment for all cells using the combined cell and nucleus feature set in (A) proximal cells and (B) distal cells and report the average balanced accuracy of pairwise classification using a 10-fold cross-validation. (C) Shows the difference in accuracies between proximal and distal cells. The insets in the top right of panels A-C show the mean result of each treatment class across all one-vs-one comparisons. (D) We tested the different feature sets using cytoplasm features only, nucleus features only and combining these two. We also tested predictions in different environments from cells proximal to the coverslip to distal from the coverslip to all cells (both proximal and distal). Here we show the results for Blebbistatin vs Nocodazole. All other treatments are shown in the Supplementary material. (E) Shows examples of our point cloud renders of 3D red blood cell shape class dataset [23]. (F) Shows the confusion matrix when classifying between these shape classes using an SVM on the learned features. (G) Shows examples of our point cloud renders of diseased and healthy 3D blood vessels from the MedMNIST v2 dataset [24]. (H) Shows the confusion matrix when classifying between an aneurysm and a healthy 3D blood vessel using an SVM on the learned features.

The accuracy levels in these pairwise classification tasks reflect the substantial overlap in the feature space of treatments described in Figure 2 C-E and the Supplementary Figure 2 A-J. Here we saw the effect of treatments does not create novel geometries but shifts the distribution of features within a population. This is likely to make binary classification at the level of single cells challenging, compared with the classification of populations of cells where changes in feature distribution are detectable.

### Learned shape features generalise to a range of 3D biological datasets

We tested the ability of automatically learned feature sets (trained on our melanoma cell dataset) to generalise on other 3D biological shape datasets. The first dataset was a library of 825 3D red blood cell shapes grouped into six and nine classes [23]. We created point cloud representations of these 3D shapes (Figure 3 E) and aligned the point clouds to a common axis (see Methods). We extracted features using the DFN trained on the WM266.4 dataset and used these features to train an SVM to predict the shape class of the 3D red blood cell (see Methods). We achieved an average balanced accuracy of 95.49% and a macro F1-score of 92.62% on an unseen test set when predicting the six classes and an average accuracy of 98.76% and a macro F1 score of 99.16% when predicting the nine classes (Figure 3 F). Previous works extracted 126 “classical” features comprising volume, roughness, and texture Gabor features and trained a random forest classifier to predict shape classes with an F1-score of 88.6% when predicting the six shape classes. Thus, our features outperformed commonly used classical features in the task of classifying 3D red blood cell shapes.

The following dataset was the VesselMNIST3D dataset, part of the MedMNIST v2 collection [24]. This dataset contains 1909 3D binary masks of healthy and aneurysm vessel segments split into training, validation and testing sets. We created point cloud representations of this dataset (Figure 3 F). We then extracted features for the training, validation, and test set using our DFN trained on the melanoma dataset. Using these features, we trained an SVM on the training data to achieve a test balanced accuracy of 83.76% and an area under the receiver operating curve (AUC) of 0.89. This is comparable to supervised methods shown in [24], which report accuracies ranging from 74.9% to 93% and AUCs ranging from 0.846 to 0.928. Most notably, our method, which extracts features in an unsupervised routine, outperformed ResNet18 + 3D (a supervised technique) in terms of accuracy and AUC (Figure 3 G).

### Interpreting the learned features

While our analyses of the latent space described by DL features suggest that they are describing aspects of cell shape, *which* aspects they describe are not immediately apparent. To interpret the learned 3D shape features, we performed principal component analysis on the DL features and correlated these with our classical, hand-crafted shape features (Figure 4 A). The first two PCs account for 42.47% of the variation in the data, with the first 11 PCs accounting for 82.24%. PC 1 correlates with many of the classical features, with major axis, surface area, and sphericity features having the highest correlation coefficients. Thus a significant degree of the variance in DL feature space can be explained by interpretable classical features. However, DL is also likely capturing properties not explained by classical feature extraction. For example, PC 3, which explains 10.85% of the variation in the data, has a maximum correlation coefficient of 0.14 with classical features (minor axis) and does not correlate highly with any classical features suggesting that the DFN may be capturing features which are not explained by our classical features.

**Figure 4:**
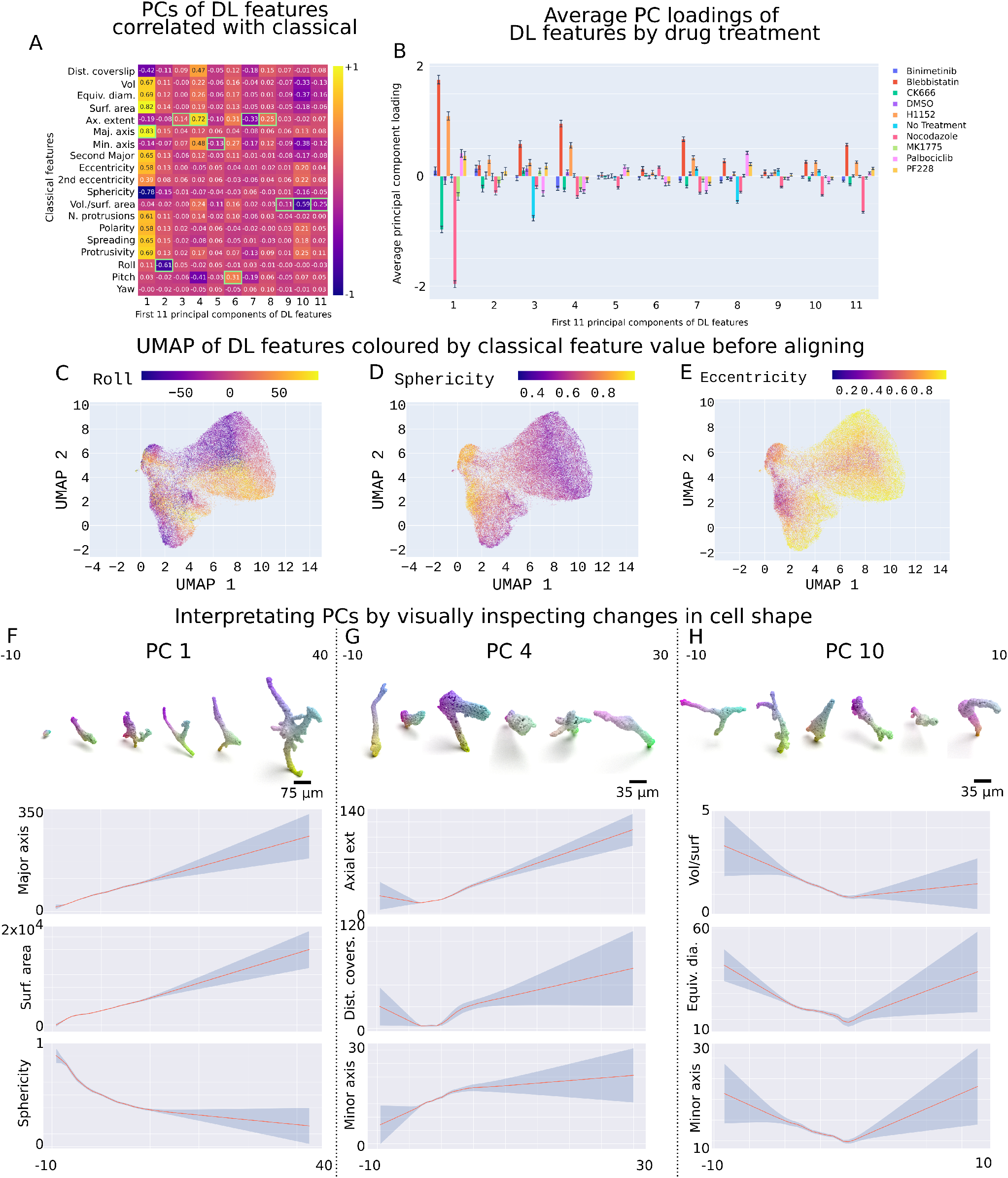
Interpreting the learned features. (A) Pearson correlation coefficients between principal components of extracted cell shape features and classical features for features. Green boxes show the classical features with the highest absolute correlation to each principal component. (B) Average PC loadings for cells treated with small molecules. Error bars indicate the standard error of the mean. (C-E) Scatter plot of UMAP embedding of learned cell shape features dataset with each point coloured by their corresponding (C) roll, (D) sphericity, and (E) eccentricity values, in order from left to right. (F-H) Rendered cell shapes were sampled from binned encodings of the respective PC value. (F) Shows this for PC 1, (G) for PC 4, and (H) shows PC 10. We show a local regression fit with a 65% confidence interval between (F) PC 3 and major axis, surface area (Surf. area), and sphericity. (G) Shows the sampled images for the range of PC 4 values. Panels show local regression fit with a 65%8confidence interval between PC 4 and axial extent, distance from the coverslip (Dist. covers.), and minor axis. (H) Shows the same for PC 10. Here, we present the local regression fits of PC 10 with volume-to-surface-area ratio (Vol/Surf), equivalent diameter (Equiv. dia.), and Minor axis.

We reasoned that if DL features were capturing meaningful biology, how cells are described in a latent space of the features should be altered by small-molecule treatments in predictable ways. Meaning DL latent space would capture changes in cell shape versus changes in the background or other non-meaningful aspects of the image. Indeed, cells treated with H1152 and Blebbistatin, which block contractility by inhibiting ROCK, and ROCK’s effector, myosin, respectively, resulted in significant similar shifts in DL PC space compared to control cells (Figure 4 B). In contrast, Nocodazole and CK666, which block protrusion (and enhance contraction), resulted in similar changes in DL PC space compared to control, which were opposite in sign compared to H1152 and Blebbistatin. Other compounds in the screen also changed DL features considerably (Figure 4 B). Thus DL features can describe similar modes of action for different molecules, i.e. regulators of contractility and protrusions are quantitatively similar in DL latent space. More so, DL features have sufficient resolution to discriminate between biological effects, such that they can detect differences between regulators of contractility and protrusion.

The ability of DL features to capture classical features was visualised by performing UMAP on the DL features and colouring each point by the corresponding classical feature value. We examined this relationship for the extracted DL features. (Figure 4 C-E show how the DL features incorporate classical features: roll, sphericity and eccentricity (see Supplementary Figure 1 for all features).

We explained the DL features that contributed the most to successful discrimination between cell treatments in different classification tasks (see Methods). Our analysis focused on the classification of Nocodazole-treated versus Blebbistatin-treated cells, as these represented two very different exemplar states in the latent space described by DL features. This approach revealed that PC 1, PC 4, and PC 10 were most important in classifying Nocodazole and Blebbistatin-treated cells (see Methods). On average, cells treated with Nocodazole had low scores of PC 1, while cells treated with Blebbistatin had high scores of PC 1. Similar to methods in [25], sampling cell shapes from the dataset with a range of values for different principal components visually confirmed the correlations with classical features (Figure 4 F-H shows this for PC 1, PC 4, and PC 10 as these were the drivers of the classification between Blebbistatin and Nocodazole. To assess the relationship between classical features and the PC of the DL features, we fit local regression curves using weighted linear least squares regression (lowess) (Figure 4 F-H). We calculated confidence intervals around the lowess fits by using bootstrapping to provide an estimate of the spread of the curve. We show the 65% confidence intervals. This shows that PC 1 of the DL features is linearly related to the major axis of elongation (Major axis) and the surface area (Surf. area) of the cell. We also see that as PC 1 increases, the sphericity of the cell decreases at a decreasing rate. The other principal components, PC 4 and PC 10, represent more complex relationships to the classical features. Taken together, we show that classical shape descriptors can often explain DL features (see Supplementary Figure 4 for the correlation between raw DL features to classical features).

### Deep embedded clustering learns shape features and classes simultaneously

While binning treatments into single, specific shape classes might be beneficial at a high level and may give insight into what the drug is doing to the cell, this binning is actually unnatural. Cell shape is a continuous variable, and the shapes of treatments may be highly overlapping. We argue that there exists a distribution of shapes that exist in the dataset and treatment conditions adjust this distribution rather than defining an exemplar shape for each condition. More so, no two cells are identical but rather similar to each other with some similarity score. We, therefore, define a 3D quantitative morphological signature (3DQMS) which is an interpretable signature of how treatments conditions are distributed across shape modes in the dataset [26], and we do this through a process called improved deep embedded clustering (IDEC) [27] (Step 2 in Figure 1).

After training the DFN autoencoder, we performed improved deep embedded clustering (IDEC). Here, the clustering layer outputs a vector of “soft labels” (or a cluster distribution) for each cell. The clustering layer and the full autoencoder are then trained further to learn features representative of the input shape and shape clusters (see Methods). The soft label is interpreted as the probability of assigning a cell (*i*), represented by its features (**z**_**i**_) of an input cell to a shape cluster (*j*) (*µ*_**j**_). For our experiments, we selected five clusters according to the elbow method on the sum of squared distances of each data point to its assigned cluster centre and the Kneedle algorithm [28].

Features from each cell were extracted using the refined encoder. Figure 5 A presents the estimated probability density of the UMAP of the DL features through a kernel density approximation after IDEC training. We sampled from the dataset for cells with the highest values of soft-label for each of the five shape modes to see modes ranging from round, elongated, to protrusive and block-shaped cells (renderings of point clouds in Figure 5 A). Calculating the average classical features for each shape cluster revealed an interpretation of the five shape clusters (Figure 5 B). After examining the shape space of drug-treated cells, we saw that Nocodazole-treated cells are made up primarily of shape cluster 1 (Figure 5 C). In contrast, Blebbistatin-treated cells explore the other four of the five shape clusters (Figure 5 D), labelled 2-5.

**Figure 5:**
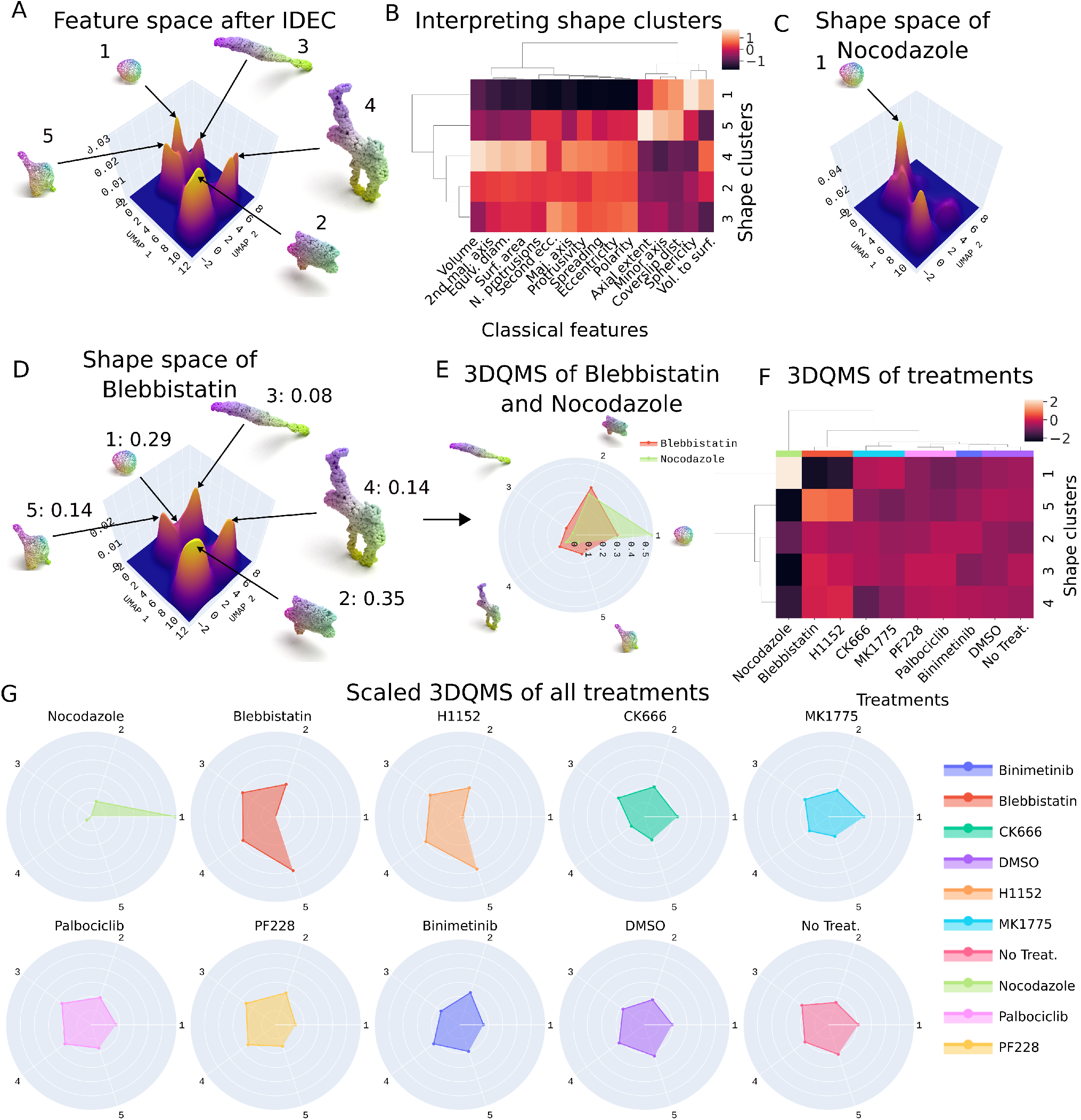
Improved deep embedded clustering learns quantitative morphological signatures for 3D shapes of cells. We added a clustering layer to a trained DFN model to learn the shape clusters that exist in the dataset. (A) Shows the kernel density approximation of the UMAP features extracted from the DFN after IDEC. The clustering layer outputs a distribution of soft labels across the five shape classes. We show point cloud renders of the five exemplar shape classes. (B) Hand-crafted measures of geometry were extracted for each cell. We described each shape cluster by these measures. (C) Shows this density approximation of the UMAP projection of the features extracted from the DFN after IDEC for Nocodazole-treated cells and (D) for Blebbistatin-treated cells. Each cell was assigned a 3DQMS, which (in this case) is a five-dimensional vector that describes how similar a cell is to each shape cluster. We also show the average similarity score of Blebbistatin-treated cells to the five shape modes. (E) Shows a radar chart of the 3DQMS for Blebbistatin and Nocodazole-treated cells. (F) We standardised and averaged the 3DQMS for each treatment and performed hierarchical clustering to reveal six classes of the treatment effects on cell shape. (G) Shows the radar chart of the average standardised 3DQMS for each small-molecule treatment.

Figure 5 D and E show examples of a 3DQMS for Nocodazole and Blebbistatin-treated cells, respectively. As shown in Figure 5, the 3DQMS for Blebbistatin is [0.29,0.35, 0.08,0.14,0.14]. A 3DQMS is a probability mass function that offers an interpretable signature that can directly observe how each condition (e.g. treatment) is spread across the common shape modes that exist in a dataset.

Next, we standardised and averaged the 3DQMS for each cell by their treatment and performed hierarchical clustering on these to find six classes of treatment effects on shape (Figure 5 F). We noted that two distinct inhibitors of myosin activation (Blebbistatin and H1152) grouped together, indicating that the 3DQMS will be a powerful tool for using cell shape to identify drugs that act on similar pathways. The other classes of treatment effects on shape included Nocodazole, which inhibits the polymerisation of microtubules, CK666 (prevents actin branching through inhibition of actin related protein (Arp) 2/3 complex) and MK1775 (WEE 1 inhibitor), PF228 (a focal adhesion kinase (FAK) inhibitor) and Palbociclib (cyclin dependent kinase (CDK) 4/6 inhibitor), Binimetinib (mitogen-activated protein kinase (MEK) inhibitor), and then the DMSO and No Treatment (control cells). The standardised 3DQMS for each treatment show how the average cell of a treatment scores across the shape modes compared to other treatments (Figure 5 G).

### A platform for genetics and drug-discovery

Next, we sought to apply our model to a comprehensive set of cell perturbations. To do this, we extracted DL shape features from WM cells that had been depleted for most human Rho guanine nucleotide exchange factors (RhoGEFs), Rho GTPase activating proteins (RhoGAPs), and Rho family GTPases. This data set comprised over 35,000 single cells across 168 different treatment conditions (RNA interference (RNAi)). OPM was used to image the 3D shape of cells.

Our DFN autoencoder without the clustering layer extracted 128 DL shape features from point cloud representations of each single cell. We normalised the shape features, and then calculated an average feature vector for each gene knockdown. Next, we calculated and clustered on a correlation matrix (Figure 6 A). We used publicly available datasets to annotate each knockdown with molecular features of the protein encoded by the gene, such as major protein domains and the specificity of each RhoGEF and RhoGAP 6 B). There existed 3 major shape clusters, with cluster 1 enriched for the depletion of RhoGAPs and clusters 2 and 3 enriched for the depletion of RhoGEFs. We interpreted these clusters by reference to classical features (Figure 6 C). Cells in cluster 1 (enriched for the depletion of RhoGAPs) tended to be larger and more protrusive. In contrast, cells in clusters 2 and 3 (enriched for the depletion of RhoGEFs) tended to be smaller and more spherical. Thus, this reveals that 3D morphology can be used to infer global protein activities, for example, whether RhoGAPs (cluster 1) or RhoGEFs (clusters 2 and 3) are ‘off’.

**Figure 6:**
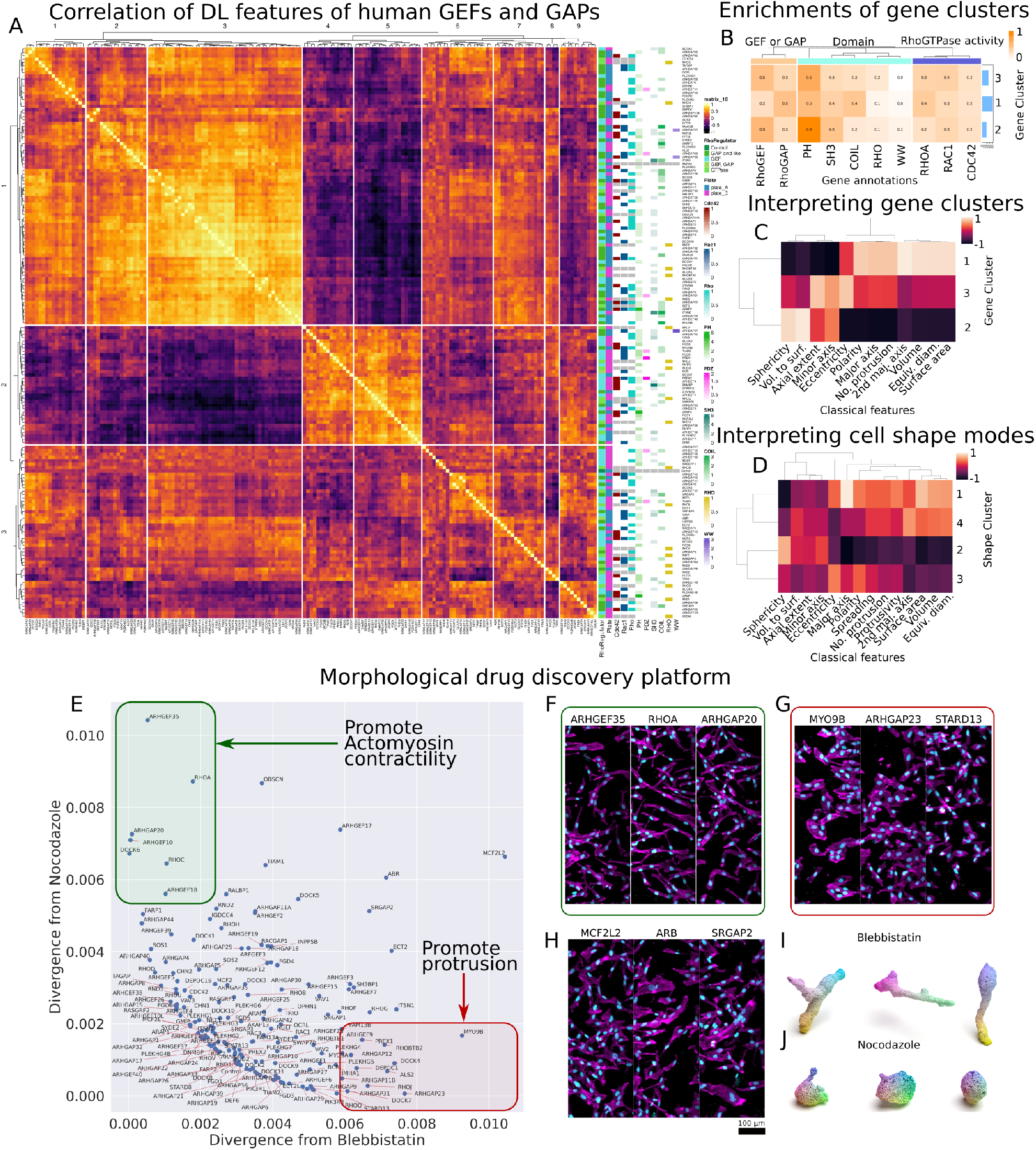
A morphological drug discovery platform. (A) Cell shape features were extracted from cells treated with RNAi. The average cell shape features of each RNAi were normalised and correlated to each other. The correlation matrix was clustered into three and nine clusters using hierarchical clustering. (B) Each GEF and GAP was annotated with its major protein domain and the GTPase it regulated. Enrichments of these annotations are shown in a cluster map. (C) The shapes of the gene clusters were interpreted using hand-crafted features. (D) The RNAi dataset and proximal cells from the treatment dataset were combined, and IDEC was performed. The four common shape modes and their hand-crafted features are shown for interpretability. Differences in the 3DQMS of the RNAi’s from Blebbistatin and Nocodazole were calculated using KL divergence. (E) Shows a scatter plot of each RNAi’s divergence from Nocodazole (x-axis) and Blebbistatin (y-axis). Proteins that are suggested to promote actomyosin contractility are highlighted in a green box, and proteins suggested to promote protrusions are highlighted in a red box. (F) Shows MIPs of ARHGEF35, RHOA, and ARHGAP20, which are highlighted in green in (E). (G) Shows MIPs of MYO9B, ARHGAP23 and STARD13 (highlighted in red in (E)). (H) Shows maximum intensity projections of MCF2L2, ARB, and SRGAP2, which are neither similar to Blebbistatin nor Nocodazole. 1(I1) Shows rendered point clouds of Blebbistatin-treated cells. (J) Shows rendered point clouds of Nocodazole-treated cells.

Furthermore, clusters 2 and 3 could be weakly distinguished based on the particular protein domains that were enriched in the targets of knockdown. With Pleckstrin homology (PH) and Src homology 3 (SH3) domains being weakly enriched in cluster 2 (Figure 6 B). In addition, cluster 3 was weakly enriched in RhoGEFs that target RhoA (Figure 6 B). Thus, 3D cell shape can reveal insight into the activation state of proteins at the domain level; that is, we can infer that PH/SH3 domain-containing proteins are off in cluster 2 but more active in clusters 1 and 3.

We then sought to determine if 3DQMS could predict the biological pathways targeted by different small-molecules by integrating data following small-molecule treatment and genetic perturbation (Figure 2 G and H). We extracted features for the cells in this combined dataset and performed k-means clustering on a range of different values for *k*_*c*_, which indicated that the optimal number of shape modes in this combined dataset was 4 (see Methods). The trained autoencoder and clustering layer revealed the four common shape modes in the dataset. Similar to methods used to generate Figure 5 E, we interpreted these shape modes by reference to classical measures of cell geometry (Figure 6 D). With the trained autoencoder and clustering layer, we extracted the 3DQMS for every cell and computed the average for each RNAi. Since this 3DQMS is a probability distribution of a treatment across shape modes, we compared each RNAi 3DQMS to the 3DQMS of Nocodazole and the 3DQMS of Blebbistatin through a KL divergence (Figure 6 E). We selected Blebbistatin and Nocodazole as they were the treatments most different from the control and each other (Figure We expected the depletion of genes that promote contractility to have a high divergence from Nocodazole. Prominent amongst this set are RhoA and RhoC, activators of the ROCK-myosin activity and contractility [29] (green box in Figure 6 E). In contrast, we expected the depletion of genes that ordinarily prevent contractility to have a high divergence from Blebbistatin. Indeed cells depleted for Myo9B, a microtubule-associated protein which normally functions to inhibit RhoGTPase-based contractility [30], are quantitatively similar to Nocodazole treatment but divergent from cells treated with Blebbistatin. Figure 6 F-H shows the maximum intensity projections of cells that have similar 3DQMS to Blebbistatin and different 3DQMS from Nocodazole (Figure 6 F), similar 3DQMS to Nocodazole and different 3DQMS from Blebbistatin (Figure 6 G), and different 3DQMS from both Nocodazole and Blebbistatin (Figure 6 H). It is evident that cells depleted for Rho-regulators that are suggested to promote actomyosin contractility are more protrusive (Figure 6 F), whereas cells depleted for genes that are suggested to promote protrusions are rounder 6 G. The cells that are both different from Nocodazole and Blebbistatin tend to be neither round nor eccentric. (Figure 6 I and J) show examples of point cloud renders of Blebbistatin, and Nocodazole treated cells, respectively. Taken together, this shows that the similarity of 3D cell shape following gene depletion or small-molecule treatment can be used to infer the molecular targets of small molecules.

## Discussion

Here, we demonstrated the utility of GDL to automatically learn 3D single-cell shape profiles, shape modes, and treatment distributions across these modes towards the goal of creating a phenotypic drug screening platform. We used these extracted features to distinguish between cancer cells treated with different small molecule treatments, 3D red blood cell shapes and diseased 3D blood vessels with accuracies greater than conventional methods and comparable with other supervised DL methods. We further defined a 3DQMS for each cell and each treatment class. This 3DQMS measures how similar the shape of each cell is to the shape modes found in the dataset and is an interpretable readout which describes a shift in cell shape distribution of treatments. We demonstrated how our methods could be used as a morphological drug discovery platform by combining our small molecule treatment dataset with a dataset of cells that had been depleted of most human RhoGEFs, RhoGAPs and RhoGTPases.

Building a deep learning model requires design choices. The first decision is the representation choice of 3D input data. Since the creation of 3D object datasets such as ModelNet [3], the two broad approaches used to represent the input data are voxels (the 3D counterpart to pixels in 2D images) and point clouds. Recently point clouds have been the dominant approach [31, 9, 32]. One reason for this is practical, and another is performance. Additionally, point cloud data is a straightforward representation of a 3D shape consisting of a 2D matrix of the positions of points along the surface. We had the choice of both voxel and point cloud representations and explored both approaches when representing human cells. For learning representations on 3D voxel grids, we utilised 3D ResNet architectures. Subsequently, we packaged our methods as a Python package called ‘`cellshape-voxel`’ for researchers to use freely. Using performance on classification tasks as a guide, we found that autoencoders using point cloud data were superior and have thus reported on these results (see supplementary material). Our choice of 2048 points was based on previous works [17, 6] and was tested and compared against three scales of sampling densities to ensure points along intricate cellular protrusions were captured while maintaining computational efficiency. Future work could explore methods of describing cells by either a few critical points or, alternatively, methods that could optimise the densities of points representing cells at different spatial resolutions. For example, a cell could be described by its centre of mass and points along its protrusions. While decreasing the detail by describing cells by much fewer points, computational efficiency would significantly increase.

A second key design decision is the selection of an autoencoder. As previously discussed, geometric deep learning is the branch of deep learning that deals with graph or manifold (unstructured) data, such as graphs created on point clouds. Geometric deep learning has predominantly been used on point cloud data to produce state-of-the-art results for classification and representation learning tasks [31, 18]. Our model incorporated edge convolution as the primary operator in the encoder part of our autoencoder, as this has proved successful in representation learning tasks [32]. FoldingNet is a folding-based decoder designed to assist representation learning on point clouds [17]. This decoder backbone has been used extensively in the literature primarily as it is simple and has shown promising results across several tasks [16, 18]. Thus, we incorporate a folding-based decoder in our model.

A third important decision is a method to group unlabelled data. This is a challenging task with methods proposed for common 2D benchmark datasets [33, 34]. To solve this problem for biological cells, we took the approach of deep embedded clustering [21]. Deep embedded clustering is a way to simultaneously learn feature representations and cluster assignments or classes. This method involves clustering with Kullback–Leibler divergence on feature representations from an autoencoder [21]. This method can easily be incorporated into any autoencoder architecture and offers an interpretable output in the form of a soft label which describes the probability of assigning data input to each cluster class. This, in turn, allows the assignment of data input to be continuous across classes rather than discrete. Cell morphology is a continuous variable with no two cells being identical in shape, but rather similar to each other or similar to exemplar shapes [35]. Therefore, describing cell shapes by a continuous similarity score to exemplar shapes offers a more natural solution than discrete binning into certain shape classes [26]. Thus, we have opted for deep embedded clustering to learn cell shapes’ 3DQMS.

A major component of our work is its accessibility to the wider biological and medical community. Packing our methods in open-source Python packages with ease of use allows further research on connecting 3D shapes to function across various domains. Our Python package offers adaptability and growth with the plan to implement new methods continuously.

## Methods

### Melanoma cell preparation

Cells used in this study were WM266.4 harbouring CAAX-EGFP (donated from the Marshall lab), with the addition of an ERK-KTR-Ruby construct (addgene #90231) and a Histone2B-iRFP670 construct (Addgene #90237).

### Collagen preparation

Collagen hydrogels were prepared to a final concentration of 2mg/mL. Briefly, hydrogel solutions of dH_2_O, 5xDMEM, HEPES (7.5 pH), and Rat Tail Collagen IV (Corning) were prepared on ice to a final collagen concentration of 2*mg/mL*. Cells were re-suspended in hydrogel solution at a concentration of 4 E4 cells per 100 *µl*, and 100 *µl* of this solution dispensed into each experimental well on a 96 well plate. After dispensing cells, the plates were incubated at 37 degrees Celsius for 1 hour, and 100 *µl* of DMEM was added to each well.

### Treatments and cell fixation

Treatments were added to cells 24 hours after seeding in collagen hydrogels. After 6 hours of treatment, cells were fixed in 4 percent paraformaldehyde for 30 minutes at room temperature. Final concentrations for treatments were: Binimetinib (2 *µM*), Palbociclib (2 *µM*), MK1775 (1 *µM*), Blebbistatin (10 *µM*), H1152 (10 *µM*), PF228 (2 *µM*), CK666 (100 *µM*), Nocodazole (1 *µM*) and DMSO 1 in 1000. Concentrations were calculated including the 100 *µl* volume of the collagen hydrogel. See supplementary materials for more information on these treatments.

### High throughput RNA interference screening on top of stiff material

RNA interference (RNAi) screens were performed in 384-well Cell Carrier plates (PerkinElmer) to which 40 *nL*/well small interfering RNA (siRNA) (20 *µM*) were plated using an Echo liquid handler (LabCyte). Prior to seeding cells, 10 *µL* of OptiMEM (Invitrogen) containing 40 *nL*/well Lipofectamine RNAiMAX (Invitrogen) was added using a Multidrop Combi Reagent Dispenser (ThermoFischer) and plates were incubated for 30 minutes at room temperature. 5000 cells per well were seeded in 20 *µL* of complete medium for all wells containing siRNA and a subset of control wells. After 48 hours, cells were fixed by adding 30 *µL* of pre-warmed 8% PFA (methanol free) (ThermoFischer), and incubated for 15 minutes at room temperature. After washing 3 times with phosphate-buffered saline (PBS), cells were permeabilised in 0.2% Triton X-100. Cells were blocked for 1 hour at room temperature with 0.5% bovine serum albumin (BSA) in PBS. After washing for three times with PBS, primary antibodies were added in 10 *µL* block solution, and plates were sealed and incubated overnight at 4^◦^C. Following three washes in PBS, secondary antibodies were added and incubated for 2 hours at room temperature. Plates were washed two times in PBS, incubated for 15 minutes with 5 *µg*/*mL* Hoechst, washed once with PBS, filled with 50 *µ* PBS, and sealed for imaging.

### Microscopy setup and image acquisition

OPM imaging was performed on a modified version of the OPM system described in [2, 19]. The primary microscope objective was a 60X/1.2NA water immersion objective, and the secondary objective was a 50X/0.95NA air objective, and the tertiary objective was a 40x/0.6NA air objective. The OPM angle was 35 degrees.

A single sCMOS camera (pco Edge) was used in Global Reset acquisition mode with 1280 × 1000 pixels. A motorised filter wheel (FW103H/M, Thorlabs) was used to switch between filters for multichannel imaging. An OPM volume was acquired for iRFP (642 *nm* excitation and 731/137 emission filter (Semrock Brightline)) before the stage returned to the start position before the acquisition of the EGFP volume (488 *nm* excitation and 550/49 emission filter (Semrock Brightline)). Finally, a collagen scattered light volume (488 *nm* illumination and no emission filter) was acquired from the same start position. The laser illumination and camera exposure time were both 4 ms. The stage velocity was 0.16 *µm*/*ms* and image acquisition was triggered every 1.4 *µm* of stage travel. For each field of view, the x-y stage covered 4000 *µm*, and three regions were imaged for each well. Before analysis, raw frames were compressed using jetraw compression software (jetraw, Dotphoton). Volumes were then de-skewed into standard *xyz* coordinates [2] and binned such that the final voxel size was 1×1×1 *µm*^3^. Image reslicing was performed using bi-linear resampling similar to the methods described in [19].

### Segmentation

The OPM image acquisition produces a parallelepiped-shaped volume due to the light-sheet angle. Before segmentation, the tips of this parallelepiped were cropped out in the y direction. This process removed sections of the volume that were not imaged over the entire axial extent. The collagen channel was viewed manually prior to segmentation, and volumes, where the collagen was not present throughout the whole volume, were rejected.

Two 3D segmentation methods were used in this paper to verify the robustness of our approach. These included Ostu’s thresholding and active contours. These methods calculate a binary mask of the region of interest with minimal user input. We used both methods to segment both cells and nuclei in 3D. Otsu’s thresholding was computed automatically for each field of view. The threshold for the nucleus was computed using Otsu’s method with a single-level (multithresh, Image 222 Processing Toolbox, MATLAB). There were cases in the data where brightly fluorescent cell membranes with dimly fluorescent protrusions existed. To include all these parts of the cell in the output segmentation, the lowest threshold found by a two-level Otsu method was used. For the active contours method, an initial guess of the segmentation mask was generated using a threshold value slightly above the intensity of the background (a value of 5). The final segmentation output was then calculated after 500 and 1000 iterations for nuclei and cells, respectively, of the active contours method using default parameters (Image Processing Toolbox, MATLAB).

Touching nuclei were separated following a two-step process. First, a Euclidean distance transform of the inverse of the mask was calculated to determine the distance of each voxel to the edge of the 3D mask (bwdist, Image Processing Toolbox, MATLAB). Next, a watershed method (Image Processing Toolbox, MATLAB) was applied to the negative of the distance-transformed image. These steps separated nuclei based on regions where the masks began to narrow. Nuclei volumes less than 50 *µm*^3^ were rejected at this stage.

Any connected components in the cell binary mask which did not contain a nucleus were rejected. Touching cells were separated using a marker-based watershed approach. Here, the nuclei are set as the low points (value of 0), the cell cytoplasm is set as intermediate points (value of 1), and the background is set as high points (value of digital infinity). This watershed method finds the halfway point between the nuclei of touching cells. Following segmentation, cells touching the volume edges were removed. Cells with volumes less than 512 *µm*^3^ were removed. The nuclei of rejected cells and cells of rejected nuclei were removed. We further tested Cellpose [36], a deep learning-based segmentation method for a wide range of microscopy images. However, after manually inspecting the results, we found that our procedure involving Ostu’s thresholding and active contour with further pre- and post-processing worked better on our dataset. This was primarily due to the cases where there were brightly fluorescent cell membranes with dimly fluorescent protrusions.

There were cases where our watershed algorithms failed and some region’s of interest (ROI) contained multiple cells. As an extra filtering step, we re-segmented the nuclei channel for each ROI. If there were more than one nucleus mask from our new segmentation that was in the same voxel locations as the cell mask from the first cell segmentation, we removed this ROI from the dataset entirely. The segmentations were manually checked for several “difficult” cells, and we are confident that our method is sufficiently robust.

### Outlier removal

The three repeated experiments produced a segmented dataset of over 65000 single cells. Similar to [19], we removed cells and nuclei that expressed low CAAX-GFP transgene, as this caused inaccurate segmentation. We automatically removed cells with a maximum intensity less than the mean of all cell maximum intensities minus the standard deviation of all cell maximum intensities.

After the above outlier removal and quality control, there was an additional round of quality control during the data analysis stage. Cells with a mean CAAX-GFP intensity of fewer than 70 units were removed from the study, and cells with a mean nuclear Histone-2B of fewer than five units were also removed from the study. The final dataset is represented in the supplementary material.

### Classical shape measurements

Similar to our previous works [19], cell and nuclei classical shape measurements were calculated by the regionprops3 function in MATLAB. Further measurements were derived from these outputs as described in [19].

### Point cloud generation

Point cloud representations of 3D objects have been used extensively in 3D shape analysis using deep learning [37, 38, 39, 40, 41]. Point clouds are arguably one of the simplest forms of shape representation and are easily obtainable from mesh objects - which are a common output of segmentation methods. We followed a two-step procedure to create point cloud representations of cell and nuclei surfaces. First, we converted the 3D binary mask to a mesh object. We used the marching cubes algorithm [42] from scikit-learn [43, 44] to extract vertices, faces and normals from the 3D binary mask. The marching cubes algorithm typically produces meshes with poor triangular quality [45]. When dealing with these meshes with poor triangular quality meshes, k-nearest neighbours graph constrictions may not give an equilateral representation of local features with irregular distances between vertices. This may adversely affect results through the topology of the 3D shape not being correct. We, therefore, used Laplacian smoothing [46] to obtain equilateral representations of local features. However, we found that this smoothing made decreased the predictive ability of our features for treatment classes (Supplementary Table 3). Trimesh [47] was used to form a mesh object from these vertices, faces and normals of cell and nuclei surfaces. Second, we sampled points from the surface of the mesh object to create a point cloud representing the shape of the cell or nuclei.

The number of points used to represent our shape data is a chosen parameter of our methods that essentially represents the resolution of our shape data. Since there is a large size discrepancy across the cell population, we are aware that using the same number of points for all cells may lead to larger cells being under-sampled and smaller cells being over-sampled. Several works on 3D shape understanding have used datasets of point cloud representations of objects ranging in scales from chairs to aeroplanes which use the same number of points for all objects [6, 31, 48, 9]. A trade-off exists between the resolution of the input point cloud and the speed of training and inference of our deep learning model. We thus needed a suitable number of points representing the input data across scales and being computationally efficient. To this end, to represent each shape, we tested three different orders of magnitude of point sampling frequencies, i.e., 1024, 2048, and 4096 points. To compare the different sampling frequencies, we visually inspected the point clouds of cells to ensure intricate cellular protrusions were being captured (Figure 1 C). Furthermore, we compared the accuracies of our classification tasks using features generated from shapes represented by all three scales of point densities to show that all sampling densities were similar in terms of classification accuracy. Ultimately, we found that 2048 points were sufficient in representing both large and small cells and were computationally efficient. We did not find cases where cell protrusions were missed with this sampling frequency. While it may be true that extremely fine details are overlooked by point cloud representations (details such as texture), global and local neighbourhood shapes are well represented by 2048 points for our datasets. Functions from PyTorch Geometric [49] were used to uniformly sample points from the surface of the mesh object. We packaged our point cloud generation pipeline in a Python package called `cellshape-helper` for anyone to use.

### Dynamic graph convolutional foldingnet autoencoder

The DFN autoencoder follows the design of the FoldingNet [17] with a dynamic graph convolutional neural network (DGCNN) as the encoder [6]. This encoder takes a 3D point cloud as input, constructs a local neighbourhood (k-nearest neighbour) graph on these points and applies convolution-like operations on the edges of connecting neighbouring point (EdgeConv). The authors [6] show translation-invariant properties of EdgeConv operations. [50] found that using the DGCNN encoder outperformed the original FoldingNet encoder regarding transfer classification accuracy, which is why we made the amendment.

The choice of *k*_*g*_ is a hyperparameter of the model architecture, and the original DGCNN paper [6] used a value of 20 for *k*_*g*_ when using 1024 points and a value of 40 when using 2048 points for the input point cloud representation of everyday objects from ModelNet40 [3]. A suitable value of *k*_*g*_ was considered through experimentation. A value of *k*_*g*_ too large would cause each point along the cell’s surface to be connected to too many neighbouring points, resulting in redundant information and a loss of local structure. Too small a value of *k*_*g*_ will result in the model degenerating into a convolutional neural network. Furthermore, larger values for *k*_*g*_ require longer training and inference times. We tested a range of different values for *k*_*g*_ to find a value that was both computationally efficient and produced the best results on our classification tasks. We know that using a k-NN graph constructed on a point cloud representation of a cell shape may bridge multiple neighbouring protrusions. When using *k* = 20, we observed that there were very few cases where protrusions were bridged together. A render of a graph constructed on a protrusive cell is presented in Supplementary Figure 5 to demonstrate this.

For our experiments, we chose *k*_*g*_ = 20 for our graph construction. We replaced the final linear layer from the original DGCNN architecture with one that outputs a desired feature vector length. The decoder takes the feature vector, **z**, as input and concatenates it with “source points”. We offer the ability to use a range of different source points. These included points sampled from a 2D plane, a Gaussian distribution or a sphere in 3D space. For the work in this paper, we use points sampled from a 2D plane. The feature vector concatenated with the source points is then passed through a series of two folding operations (defined in [17]) to output a reconstructed point cloud. Our model outputs 2025 points. The number of reconstructed points does not need to be the same as the input number of points. The Chamfer distance (CD) [51] is commonly used to compare two PCs. We used the extended CD presented in [17] as our reconstruction error between input point cloud *S* and reconstructed point cloud *Ŝ*. The CD is defined in Equation 1.

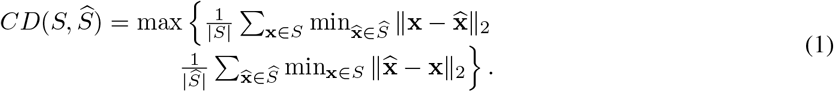

Initially, the DFN was trained on the ShapeNet dataset for 250 epochs. Then we continued training on our point cloud representations of both cells and nuclei for another 250 epochs using Adam optimiser with *e*^−6^ weight decay. We used a batch size of 16 with an initial learning rate of 0.0001 and an exponential learning rate decay scheduler. The DFN model was set to extract 128 features from each point cloud. All algorithms were implemented in PyTorch. We used code from [50, 18] and packaged our DFN into a Python package called `cellshape-cloud`.

### Deep embedded clustering

Deep embedded clustering (DEC) [21] is a specialised clustering technique that simultaneously learns feature representations and cluster assignments using autoencoders. Our implementation of DEC learns autoencoder parameters *θ* which map the 3D shapes into embedded feature space **Z** as well as *k*_*c*_ cluster centres 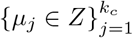 of the embedded feature space **Z**. This is done in two phases:

1. parameter initialisation with an autoencoder (through our DFN model) and
2. parameter optimisation through simultaneous autoencoder reconstruction and minimisation of the Kullback-Leibler (KL) divergence between a target distribution and a distribution of soft labels.

The second step is done by adding a clustering layer on the features to refine them by learning features that are optimised to represent the 3D shape as best as possible and grouping similar and separating dissimilar objects. This part of the model works by initialising cluster centres using the k-means clustering algorithm on the embedded feature space outputted from a pre-trained autoencoder in step 1. These cluster centres are kept as trainable parameters. The clustering layer then assigns soft labels (*q*_*ij*_) to each input by mapping features to clusters based on the Student’s t-distribution as a kernel that represents the similarity between a feature vector (**z**_**i**_) and a cluster centre (*µ*_*j*_):

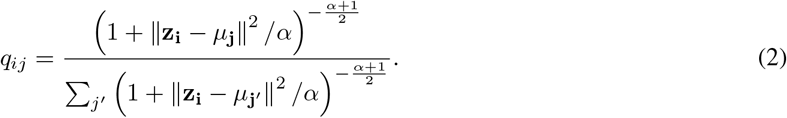

This can be interpreted as the probability of assigning input *i* to cluster *j*, hence why this is a soft assignment. Soft assignments with high probabilities are considered trustworthy; thus, DEC designs a target distribution, which raises this to the second power to place more emphasis on these confident assignments. Following DEC, we define the target distribution as follows:

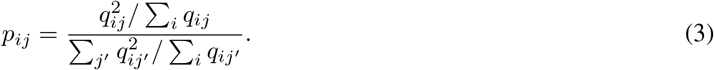

A clustering loss is defined as the Kullback–Leibler divergence between **P** and **Q**:

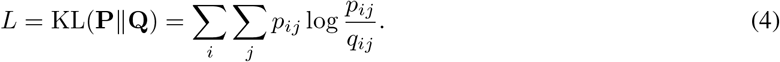

The features at cluster centres become representations of ‘example’ or ‘template’ shapes of each shape class. [21] proposed training an autoencoder in the first phase (parameter initialisation) and then abandoning the decoder in the second phase to only fine-tune the encoder through the clustering loss alone. Variants of DEC since then have shown that this fine-tuning may distort the embedded space and weaken its representativeness of the input [52, 53]. Thus, we followed the procedure in [27] and have added the decoder back to the second phase of training and optimised both the reconstruction loss and the clustering loss together with a final loss defined as:

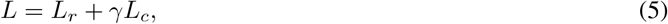

where *γ* ≥ 0 defines the magnitude that the clustering loss adds to the final loss. For all experiments, we used *γ* = 100.

Similar to methods in [27], during training of the deep embedded clustering, we updated the target distribution *p* at regular step intervals, *T*. As the target distribution constantly updates, the algorithm will not converge; thus, we needed another metric to determine when to stop training. After *T* steps, we calculated the new target distribution and divergence, *δ*, from the previous update. This divergence is the proportion of data points or cells that have changed which cluster they belong to. We set a divergence tolerance, *δ*_*tol*_, such that we stop training when *δ < δ*_*tol*_. For all our experiments, we used *δ* = 0.001

### Predicting small molecule treatments, red blood cell shapes, and healthy blood vessels

We used our extracted features to train a support vector machine (SVM). For all experiments, we used one-versus-one SVM classifiers with a radial basis function kernel and an L2 regularisation parameter (C = 5), balanced class weights, and intercept scaling during training. We used scikit-learn for these methods [43, 44]. For our drug-treated dataset we performed 10-fold cross-validation for each experiment and report the mean balanced accuracy and for the 3D red blood cell dataset, we used a 80/20 train and test set and the results on the test set. For the VesselMNIST3D dataset, we used the specified training and validation set to train our model and report on the unseen test set. For all experiments„ we report the average class accuracy to account for class imbalances.

### Feature importance

Extreme gradient boosting (XGBoost) is a machine learning algorithm which uses gradient-boosted decision trees for classification, and regression tasks [54]. A benefit of using gradient boosting is that it is relatively simple to explore which features are important for the task at hand. Importance provides a score that indicates how valuable each feature was in constructing the boosted trees in the model. The more a feature is utilised to classify, the higher its importance. This is based on the number of times a feature is used in a tree. We used this to interpret which features were important for our classification tasks. We trained an XGBoost classification model to classify Blebbistatin and Nocodazole-treated cells based on the principal components of the cell shape features extracted using the DFN without the addition of the clustering layer. For methods involving XGBoost, we used the open-source Python package ‘xgboost’ [54].

### Aligning 3D shapes to a common axis

As a step towards rotational invariance, we converted the point cloud representations of 3D red blood cells to their PCA-based canonical poses. PCA calculates three orthogonal bases or principal axis of point cloud data. This enables us to align original point clouds to the world Cartesian plane [55]. We briefly discuss how the canonical pose is calculated by following directly from [55]. We performed PCA on a given point cloud, **S** ∈ ℝ^*n*×3^, by:

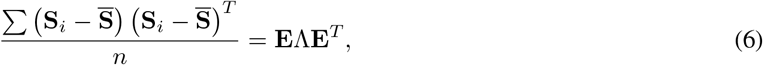

where **S**_*i*_ ∈ ℝ^3^ is the *i*^*th*^ point of **S**, 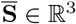 is the mean of **S, E** is the eigenvector matrix composed of eigenvectors (**e**_1_, **e**_2_, **e**_3_) (principal axes), and Λ = *diag* (*λ*_1_, *λ*_2_, *λ*_3_) are the corresponding eigenvalues. By aligning the principal axes to the three axes of the world coordinate, we obtain the canonical pose as **S**_*can*_ = **SE**.

[55] proved the rotational invariant property of **S**_*can*_. We calculated the canonical poses for each point cloud representation of the only the 3D red blood cells in used this as input to our DFN.

### Normalisation cells to the plate control

We normalised the shape features to the features of each plate’s control cells. This was done by calculating the mean and standard deviation of the shape features for the control cells of each plate. We subtracted the plate control mean for every single cell and divided this result by the plate control standard deviation.

### Selecting the number of clusters

We used the elbow method on the sum of squared distances of each data point to its assigned cluster centre and the Kneedle algorithm [28]. This was implemented using the `yellowbrick` Python package [56].

### Combining the RNAi dataset with the drug-treated dataset

We saw that cell shape depends on the environment, and the shape space of cells proximal to the coverslip differs from that of cells distal to the coverslip (Figure 2 G and H). Since our RNAi dataset is of cells on the coverslip (2D cell culture environment imaged in 3D), we combined this dataset with only the proximal cells from the drug-treated dataset.

### 3D rendering

3D renders of intensity images are generated as a 3D projection with trilinear interpolation by using the volume viewer Fiji plugin [57]. 3D renders of cell masks are presented as 3D surface representations using isosurfaces in napari [58]. Point cloud renderings of 3D cell shapes were done using Mitsuba2 [59] and adapted scripts from [60].

## Data availability

Our data are available at https://sandbox.zenodo.org/record/1170226#.ZAs8w4DP3JU. The raw data of the regions of interest and the cell and nuclei mask data will be deposited in an online repository.

## Code availability

The described software is available as a Python package which can be installed through pip, and the source code is available at https://github.com/Sentinal4D/cellshape. This is made up of four separate sub-packages for point cloud generation (cellshape-helper), graph-based representation learning (cellshape-cloud), deep embedded clustering (cellshape-cluster), as well as deep learning methods on 3D voxel input data (cellshape-voxel). Our data collection code is available at https://github.com/ImperialCollegeLondon/cellshape-collection. The data collection code includes scripts for image acquisition by OPM and our 3D cell and nuclei segmentation methods.

## Footnotes

4 https://github.com/Sentinal4D/cellshape

## References

[1] C. Dunsby. Optically sectioned imaging by oblique plane microscopy. Optics Express, 16(25):20306–20316, December 2008.

[2] Vincent Maioli, George Chennell, Hugh Sparks, Tobia Lana, Sunil Kumar, David Carling, Alessandro Sardini, and Chris Dunsby. Time-lapse 3-d measurements of a glucose biosensor in multicellular spheroids by light sheet fluorescence microscopy in commercial 96-well plates. Scientific Reports, 6(1):37777, Nov 2016.

[3] Zhirong Wu, Shuran Song, Aditya Khosla, Fisher Yu, Linguang Zhang, Xiaoou Tang, and Jianxiong Xiao. 3d shapenets: A deep representation for volumetric shapes, 2014.

[4] Angel X. Chang, Thomas Funkhouser, Leonidas Guibas, Pat Hanrahan, Qixing Huang, Zimo Li, Silvio Savarese, Manolis Savva, Shuran Song, Hao Su, Jianxiong Xiao, Li Yi, and Fisher Yu. ShapeNet: An Information-Rich 3D Model Repository. Technical Report arXiv:1512.03012 [cs.GR], Stanford University — Princeton University — Toyota Technological Institute at Chicago, 2015.

[5] Mikaela Angelina Uy, Quang-Hieu Pham, Binh-Son Hua, Duc Thanh Nguyen, and Sai-Kit Yeung. Revisiting point cloud classification: A new benchmark dataset and classification model on real-world data. In International Conference on Computer Vision (ICCV), 2019.

[6] Yue Wang, Yongbin Sun, Ziwei Liu, Sanjay E. Sarma, Michael M. Bronstein, and Justin M. Solomon. Dynamic graph cnn for learning on point clouds. ACM Transactions on Graphics (TOG), 2019.

[7] Cheng Zhang, Haocheng Wan, Shengqiang Liu, Xinyi Shen, and Zizhao Wu. Pvt: Point-voxel transformer for 3d deep learning. arXiv preprint arXiv:2108.06076, 2021.

[8] Tiange Xiang, Chaoyi Zhang, Yang Song, Jianhui Yu, and Weidong Cai. Walk in the cloud: Learning curves for point clouds shape analysis. In Proceedings of the IEEE/CVF International Conference on Computer Vision (ICCV), pages 915–924, October 2021.

[9] Xu Ma, Can Qin, Haoxuan You, Haoxi Ran, and Yun Fu. Rethinking network design and local geometry in point cloud: A simple residual MLP framework. In International Conference on Learning Representations, 2022.

[10] Jiajun Wu, Chengkai Zhang, Tianfan Xue, William T Freeman, and Joshua B Tenenbaum. Learning a probabilistic latent space of object shapes via 3d generative-adversarial modeling. In Advances in Neural Information Processing Systems, pages 82–90, 2016.

[11] Panos Achlioptas, Olga Diamanti, Ioannis Mitliagkas, and Leonidas J Guibas. Learning representations and generative models for 3d point clouds. arXiv preprint arXiv:1707.02392, 2017.

[12] Yongheng Zhao, Tolga Birdal, Haowen Deng, and Federico Tombari. 3d point capsule networks. In Conference on Computer Vision and Pattern Recognition (CVPR), 2019.

[13] Peng-Shuai Wang, Yu-Qi Yang, Qian-Fang Zou, Zhirong Wu, Yang Liu, and Xin Tong. Unsupervised 3D learning for shape analysis via multiresolution instance discrimination. In AAAI Conference on Artificial Intelligence (AAAI), 2021.

[14] Yi Shi, Mengchen Xu, Shuaihang Yuan, and Yi Fang. Unsupervised deep shape descriptor with point distribution learning. In The IEEE/CVF Conference on Computer Vision and Pattern Recognition (CVPR), June 2020.

[15] Michael M. Bronstein, Joan Bruna, Yann LeCun, Arthur Szlam, and Pierre Vandergheynst. Geometric deep learning: Going beyond euclidean data. IEEE Signal Processing Magazine, 34(4):18–42, 2017.

[16] Haowen Deng, Tolga Birdal, and Slobodan Ilic. Ppf-foldnet: Unsupervised learning of rotation invariant 3d local descriptors. In The European Conference on Computer Vision (ECCV), September 2018.

[17] Yaoqing Yang, Chen Feng, Yiru Shen, and Dong Tian. Foldingnet: Point cloud auto-encoder via deep grid deformation, 2018.

[18] Jiahao Pang, Duanshun Li, and Dong Tian. Tearingnet: Point cloud autoencoder to learn topology-friendly representations. In IEEE Conference on Computer Vision and Pattern Recognition (CVPR), 2021.

[19] L. G. Dent, N. Curry, H. Sparks, V. Bousgouni, V. Maioli, S. Kumar, I. Munro, C. Dunsby, and C. Bakal. Environmentally dependent and independent control of cell shape determination by rho gtpase regulators in melanoma. bioRxiv, 2021.

[20] William E. Lorensen and Harvey E. Cline. Marching cubes: A high resolution 3d surface construction algorithm. In Proceedings of the 14th Annual Conference on Computer Graphics and Interactive Techniques, SIGGRAPH ‘87, page 163–169, New York, NY, USA, 1987. Association for Computing Machinery.

[21] Junyuan Xie, Ross Girshick, and Ali Farhadi. Unsupervised deep embedding for clustering analysis. In Maria Florina Balcan and Kilian Q. Weinberger, editors, Proceedings of The 33rd International Conference on Machine Learning, volume 48 of Proceedings of Machine Learning Research, pages 478–487, New York, New York, USA, 20–22 Jun 2016. PMLR.

[22] Leland McInnes, John Healy, and James Melville. Umap: Uniform manifold approximation and projection for dimension reduction, 2018.

[23] Greta Simionato, Konrad Hinkelmann, Revaz Chachanidze, Paola Bianchi, Elisa Fermo, Richard van Wijk, Marc Leonetti, Christian Wagner, Lars Kaestner, and Stephan Quint. Red blood cell phenotyping from 3d confocal images using artificial neural networks. PLOS Computational Biology, 17(5):1–17, 05 2021.

[24] Jiancheng Yang, Rui Shi, Donglai Wei, Zequan Liu, Lin Zhao, Bilian Ke, Hanspeter Pfister, and Bingbing Ni. Medmnist v2 – a large-scale lightweight benchmark for 2d and 3d biomedical image classification. arXiv, 2021.

[25] Christopher J. Soelistyo, Giulia Vallardi, Guillaume Charras, and Alan R. Lowe. Learning biophysical determinants of cell fate with deep neural networks. Nature Machine Intelligence, 2022.

[26] Chris Bakal, John Aach, George Church, and Norbert Perrimon. Quantitative morphological signatures define local signaling networks regulating cell morphology. Science (New York, N.Y.), 316(5832):1753–1756, June 2007. Number: 5832.

[27] Xifeng Guo, Long Gao, Xinwang Liu, and Jianping Yin. Improved deep embedded clustering with local structure preservation. In Proceedings of the Twenty-Sixth International Joint Conference on Artificial Intelligence, IJCAI-17, pages 1753–1759, 2017.

[28] Ville Satopaa, Jeannie Albrecht, David Irwin, and Barath Raghavan. Finding a “kneedle” in a haystack: Detecting knee points in system behavior. In 2011 31st International Conference on Distributed Computing Systems Workshops, pages 166–171, 2011.

[29] Miguel Vicente-Manzanares, Xuefei Ma, Robert S Adelstein, and Alan Rick Horwitz. Non-muscle myosin ii takes centre stage in cell adhesion and migration. Nature reviews. Molecular cell biology, 10(11):778–790, 11 2009.

[30] Peter J. Hanley, Yan Xu, Moritz Kronlage, Kay Grobe Peter Schön, Jian Song, Lydia Sorokin, Albrecht Schwab, and Martin Bähler. Motorized rhogap myosin ixb (myo9b) controls cell shape and motility. Proceedings of the National Academy of Sciences, 107(27):12145–12150, 2010.

[31] Seyed Saber Mohammadi, Yiming Wang, and Alessio Del Bue. Pointview-gcn: 3d shape classification with multi-view point clouds. In 2021 IEEE International Conference on Image Processing (ICIP), pages 3103–3107. IEEE, 2021.

[32] Siming Yan, Zhenpei Yang, Haoxiang Li, Li Guan, Hao Kang, Gang Hua, and Qixing Huang. Implicit autoencoder for point cloud self-supervised representation learning. arXiv preprint arXiv:2201.00785, 2022.

[33] Mathilde Caron, Piotr Bojanowski, Armand Joulin, and Matthijs Douze. Deep clustering for unsupervised learning of visual features. In European Conference on Computer Vision, 2018.

[34] Wouter Van Gansbeke, Simon Vandenhende, Stamatios Georgoulis, Marc Proesmans, and Luc Van Gool. Scan: Learning to classify images without labels, 2020.

[35] Matt De Vries and Chris Bakal. What do machines see? Utilizing artificial intelligence to explore cell biology. The Biochemist, 43(5):48–52, 09 2021.

[36] Carsen Stringer, Tim Wang, Michalis Michaelos, and Marius Pachitariu. Cellpose: a generalist algorithm for cellular segmentation. Nature Methods, 18(1):100–106, Jan 2021.

[37] Seyed Saber Mohammadi, Yiming Wang, and Alessio Del Bue. Pointview-gcn: 3d shape classification with multi-view point clouds. In 2021 IEEE International Conference on Image Processing (ICIP), pages 3103–3107, 2021.

[38] Haoxi Ran, Jun Liu, and Chengjie Wang. Surface representation for point clouds. In Proceedings of the IEEE/CVF Conference on Computer Vision and Pattern Recognition (CVPR), pages 18942–18952, June 2022.

[39] Antonio Montanaro, Diego Valsesia, and Enrico Magli. Rethinking the compositionality of point clouds through regularization in the hyperbolic space, 2022.

[40] Hengshuang Zhao, Li Jiang, Jiaya Jia, Philip H.S. Torr, and Vladlen Koltun. Point transformer. In Proceedings of the IEEE/CVF International Conference on Computer Vision (ICCV), pages 16259–16268, October 2021.

[41] Xumin Yu, Lulu Tang, Yongming Rao, Tiejun Huang, Jie Zhou, and Jiwen Lu. Point-bert: Pre-training 3d point cloud transformers with masked point modeling. In Proceedings of the IEEE/CVF Conference on Computer Vision and Pattern Recognition (CVPR), pages 19313–19322, June 2022.

[42] William E. Lorensen and Harvey E. Cline. Marching cubes: A high resolution 3d surface construction algorithm. SIGGRAPH Comput. Graph., 21(4):163–169, August 1987.

[43] F. Pedregosa, G. Varoquaux, A. Gramfort, V. Michel, B. Thirion, O. Grisel, M. Blondel, P. Prettenhofer, R. Weiss, V. Dubourg, J. Vanderplas, A. Passos, D. Cournapeau, M. Brucher, M. Perrot, and E. Duchesnay. Scikit-learn: Machine learning in Python. Journal of Machine Learning Research, 12:2825–2830, 2011.

[44] Lars Buitinck, Gilles Louppe, Mathieu Blondel, Fabian Pedregosa, Andreas Mueller, Olivier Grisel, Vlad Niculae, Peter Prettenhofer, Alexandre Gramfort, Jaques Grobler, Robert Layton, Jake VanderPlas, Arnaud Joly, Brian Holt, and Gaël Varoquaux. API design for machine learning software: experiences from the scikit-learn project. In ECML PKDD Workshop: Languages for Data Mining and Machine Learning, pages 108–122, 2013.

[45] Lis Custodio, Sinesio Pesco, and Claudio Silva. An extended triangulation to the marching cubes 33 algorithm. Journal of the Brazilian Computer Society, 25(1):6, Jun 2019.

[46] J. Vollmer, R. Mencl, and H. Müller. Improved laplacian smoothing of noisy surface meshes. Computer Graphics Forum, 18(3):131–138, 1999.

[47] Dawson-Haggerty et al. trimesh.

[48] Benjamin Eckart, Wentao Yuan, Chao Liu, and Jan Kautz. Self-supervised learning on 3d point clouds by learning discrete generative models. In 2021 IEEE/CVF Conference on Computer Vision and Pattern Recognition (CVPR), pages 8244–8253, 2021.

[49] Matthias Fey and Jan E. Lenssen. Fast graph representation learning with PyTorch Geometric. In ICLR Workshop on Representation Learning on Graphs and Manifolds, 2019.

[50] An Tao. Unsupervised point cloud reconstruction for classific feature learning. https://github.com/AnTao97/UnsupervisedPointCloudReconstruction, 2020.

[51] Haoqiang Fan, Hao Su, and Leonidas Guibas. A point set generation network for 3d object reconstruction from a single image. In 2017 IEEE Conference on Computer Vision and Pattern Recognition (CVPR), pages 2463–2471, 2017.

[52] Xifeng Guo, Xinwang Liu, En Zhu, and Jianping Yin. Deep clustering with convolutional autoencoders. In Derong Liu, Shengli Xie, Yuanqing Li, Dongbin Zhao, and El-Sayed M. El-Alfy, editors, Neural Information Processing, pages 373–382, Cham, 2017. Springer International Publishing.

[53] Xifeng Guo, En Zhu, Xinwang Liu, and Jianping Yin. Deep embedded clustering with data augmentation, 14–16 Nov 2018.

[54] Tianqi Chen and Carlos Guestrin. XGBoost: A scalable tree boosting system. In Proceedings of the 22nd ACM SIGKDD International Conference on Knowledge Discovery and Data Mining, KDD ‘16, pages 785–794, New York, NY, USA, 2016. ACM.

[55] Feiran Li, Kent Fujiwara, Fumio Okura, and Yasuyuki Matsushita. A closer look at rotation-invariant deep point cloud analysis. In Proceedings of the IEEE/CVF International Conference on Computer Vision (ICCV), pages 16218–16227, October 2021.

[56] Benjamin Bengfort, Rebecca Bilbro, Nathan Danielsen, Larry Gray, Kristen McIntyre, Prema Roman, Zijie Poh, et al. Yellowbrick, 2018.

[57] Johannes Schindelin, Ignacio Arganda-Carreras, Erwin Frise, Verena Kaynig, Mark Longair, Tobias Pietzsch, Stephan Preibisch, Curtis Rueden, Stephan Saalfeld, Benjamin Schmid, Jean-Yves Tinevez, Daniel James White, Volker Hartenstein, Kevin Eliceiri, Pavel Tomancak, and Albert Cardona. Fiji: an open-source platform for biological-image analysis. Nature Methods, 9(7):676–682, Jul 2012.

[58] Nicholas Sofroniew, Talley Lambert, Kira Evans, Juan Nunez-Iglesias, Grzegorz Bokota, Matthias Bussonnier, Gonzalo Peña-Castellanos, Philip Winston, Kevin Yamauchi, Draga Doncila Pop, Pam, Ziyang Liu, Ahmet Can Solak alisterburt, Genevieve Buckley, Andy Sweet, Lorenzo Gaifas, Gregory Lee, Jaime Rodríguez-Guerra, Nathan Clack, Jordão Bragantini, Lukasz Migas, Volker Hilsenstein, Melissa Weber Mendonça, Robert Haase Hector, Jeremy Freeman, Peter Boone, Alan R Lowe, and Christoph Gohlke. napari/napari: 0.4.13rc0, January 2022.

[59] Merlin Nimier-David, Delio Vicini, Tizian Zeltner, and Wenzel Jakob. Mitsuba 2: A retargetable forward and inverse renderer. ACM Trans. Graph., 38(6), nov 2019.

[60] Tolga Birdal. Mitsuba2PointCloudRenderer. https://github.com/tolgabirdal/Mitsuba2PointCloudRenderer, 2020.

