## supplementary material for "3D single-cell shape analysis using geometric deep learning"

A PREPRINT

### 1 Supplementary tables

| Drug | Intended Target | Dose ($\mu M$) |
|---|---|---|
| Binimetinib | MEK 1/2 | 2 |
| Blebbistatin | NMM type-2 | 10 |
| CK666 | ARP 2/3 | 100 |
| H1152 | Rho-Kinase (ROCK) | 10 |
| Nocodazole | Microtubules | 1 |
| MK1775 | WEE 1 | 1 |
| Palbociclib | CDK 4/6 | 2 |
| PF228 | FAK (PTK2) | 2 |

Table 1: The small molecules, their intended targets and the doses used in this study.

| Treatment | Proximal | Distal | All |
|---|---|---|---|
| Binimetinib | 3693 | 3199 | 6892 |
| Blebbistatin | 2962 | 3134 | 6096 |
| CK666 | 3207 | 3356 | 6563 |
| DMSO | 3408 | 3156 | 6564 |
| H1152 | 2922 | 3156 | 6078 |
| MK1775 | 3689 | 3249 | 6938 |
| No treatment | 3072 | 3309 | 6381 |
| Nocodazole | 3352 | 3440 | 6792 |
| PF228 | 3538 | 2985 | 6523 |
| Palbociclib | 3645 | 2953 | 6598 |
| Total | 33537 | 31963 | 65500 |

Table 2: Number of cells in each treatment and environment group.

| Feature Extraction Method | Cell | Nuc | Both |
|---|---|---|---|
| Classical | 82.9 | 63.4 | 83.3 |
| 3D Conv AE | 83.1 | 71.6 | 84.3 |
| DFN smoothed | 75.74 | 64.2 | 78.11 |
| DFN unsmoothed | **83.4** | **77.2** | **86.2** |

Table 3: A comparison of features on predicting Blebbistatin from Nocodazole-treated cells.

### 2 Supplementary figures

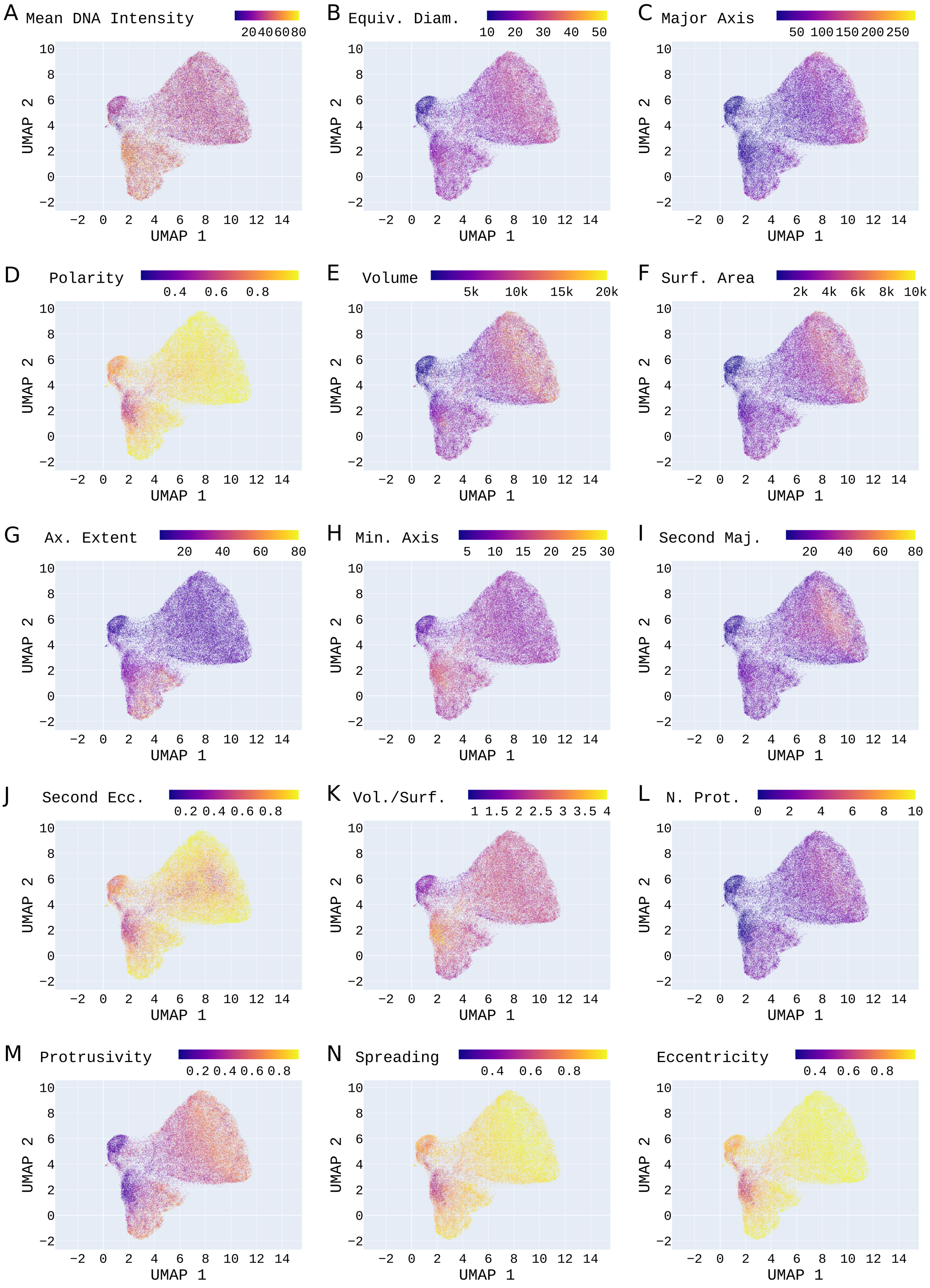


Figure 1: **Visualising patterns of classical features over the learned feature landscape.** Each panel represents a different classical feature. (A) Mean DNA intensity. (B) Equivalent diameter. (C) Major axis. (D) Polarity. (E) Volume. (F) Surface area. (G) Axial extent. (H) Minor axis. (I) Second major axis. (J) Eccentricity. (K) Volume to surface area ratio. (L) The number of protrusions. (M) Protrusivity. (N) Spreading. (O) Second eccentricity.

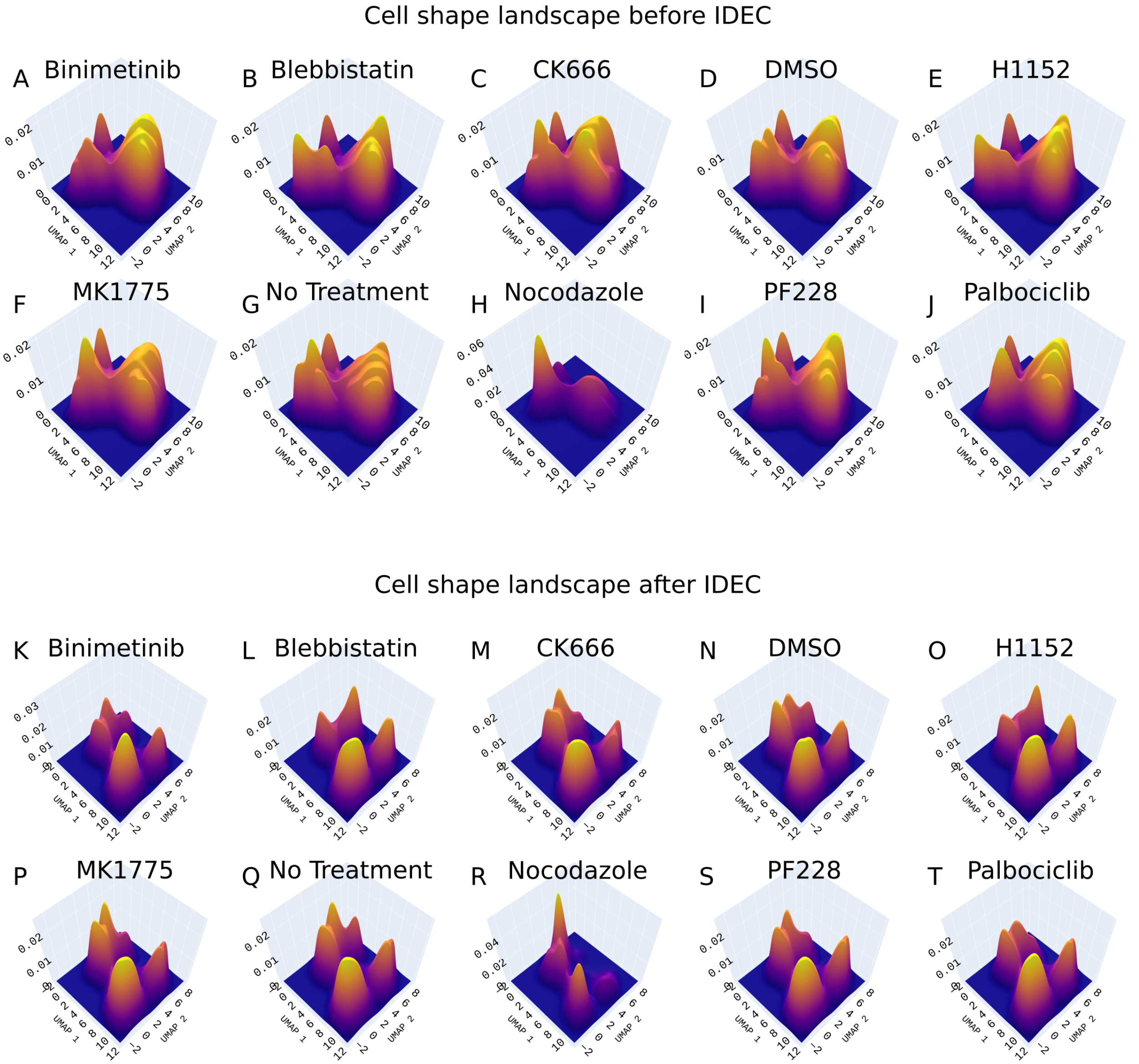


Figure 2: **Cell shape landscape of all drugs before (A-J) and after (K-T) IDEC.** Kernel density approximation of the UMAP of cell shape features of cells treated with (A) Binimetinib, (B) Blebbistatin, (C) CK666, (D) DMSO, (E) H1152, (F) MK1775, (G) No Treatment, (H) Nocodazole, (I) PF228, and (J) Palbociclib before IDEC. Kernel density approximation of the UMAP of cell shape features of cells treated with (K) Binimetinib, (L) Blebbistatin, (M) CK666, (N) DMSO, (O) H1152, (P) MK1775, (Q) No Treatment, (R) Nocodazole, (S) PF228, and (T) Palbociclib after IDEC.

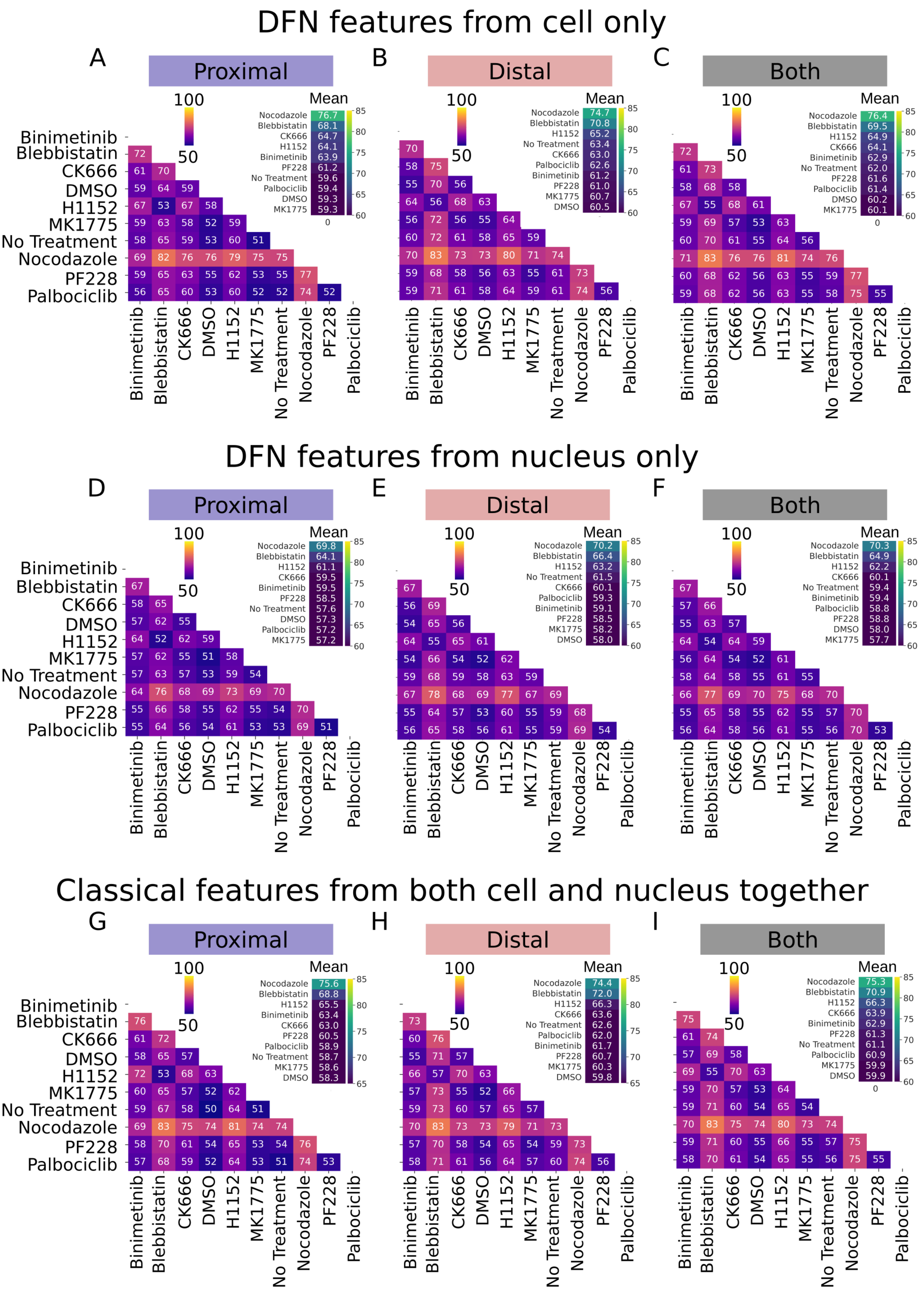


Figure 3: **Predicting drug treatment using different feature sets.** (A-C) Shows the pairwise predictions using DFN features extracted from the cell cytoplasm only. We show the different environments for this with (A) showing the proximal environment, (B) showing the distal environment and (C) showing all cells in all environments. (D-F) Show this for DFN features extracted from the nucleus. (G-I) Shows this for classical features extracted from both the nucleus and the cytoplasm.

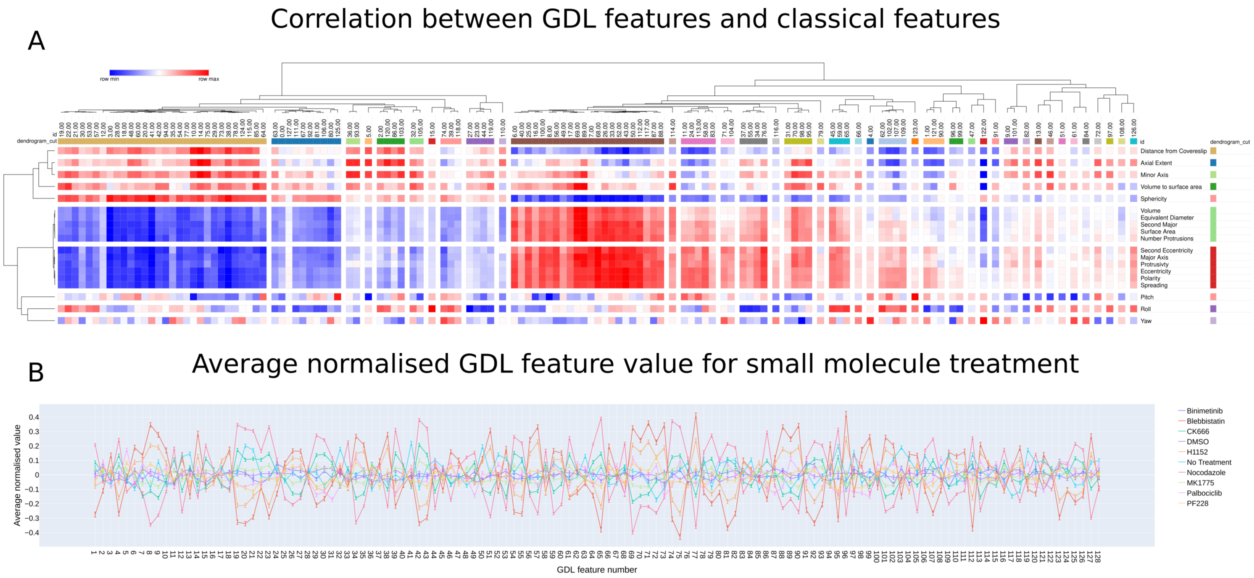


Figure 4: **Explaining GDL features with respect to classical features and small-molecule treatment.** (A) The correlation between features extracted from our DFN autoencoder to classical features of cell geometry. (B) The average normalised GDL feature values for each small molecule treatment.

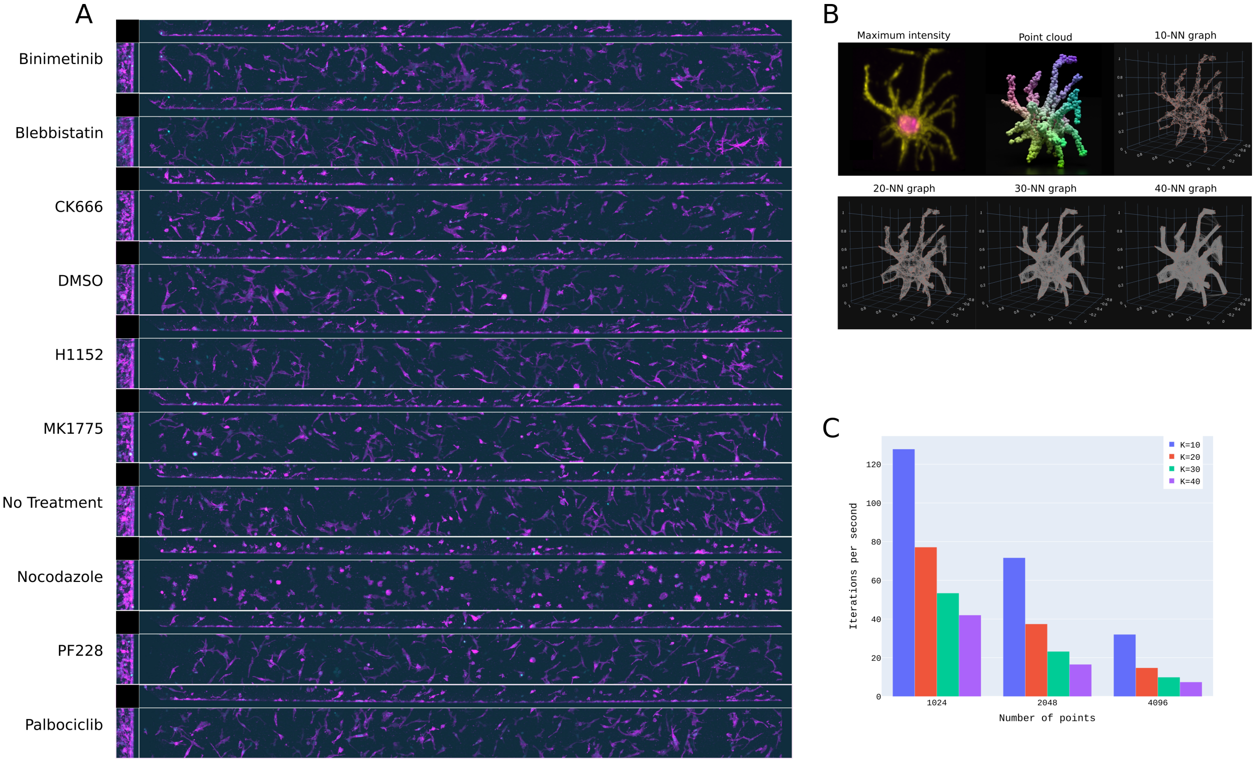


Figure 5: (A) Exampled field of views of drug-treated WM226.4 melanoma cells embedded in a collagen matrix. (B) An example of $k$-nn graphs constructed on a point cloud representation of a protrusive cell, with different values of $k$. (C) How the value of $k_g$ and number of points affects the number of iterations per second on a Nvidia Quadro RTX 6000 GPU.
